# Blood and site of disease inflammatory profiles differ in HIV-1-infected pericardial tuberculosis patients

**DOI:** 10.1101/2022.10.21.513232

**Authors:** Hygon Mutavhatsindi, Elsa Du Bruyn, Sheena Ruzive, Patrick Howlett, Alan Sher, Katrin D. Mayer-Barber, Daniel L. Barber, Mpiko Ntsekhe, Robert J. Wilkinson, Catherine Riou

## Abstract

**Objectives:** To better understand the pathogenesis of pericardial tuberculosis (PCTB), we sought to characterize the systemic inflammatory profile in HIV-1-infected participants with latent TB infection (LTBI), pulmonary TB (PTB) and PCTB.

**Methods:** Using Luminex, we measured 39 analytes in pericardial fluid (PCF) and paired plasma from 18 PCTB participants, and plasma from 16 LTBI and 20 PTB. Follow-up plasma samples were also obtained from PTB and PCTB participants. HLA-DR expression on Mtb-specific CD4 T cells was measured in baseline samples using flow cytometry.

**Results:** Assessment of the overall systemic inflammatory profile by principal component analysis showed that the inflammatory profile of active TB participants was distinct from the LTBI group, while PTB patients could not be distinguished from those with PCTB. In the LTBI group, 12 analytes showed a positive association with plasma HIV-1 viral load, and most of these associations were lost in the diseased groups. When comparing the inflammatory profile between PCF and paired blood, we found that the concentrations of most analytes (24/39) were elevated at site of disease. However, the inflammatory profile in PCF partially mirrored inflammatory events in the blood. After TB treatment completion, the overall plasma inflammatory profile reverted to those observed in the LTBI group. Lastly, HLA-DR expression showed the best performance for TB diagnosis compared to previously described biosignatures built from soluble markers.

**Conclusion:** Our results describe the inflammatory profile associated with PTB and PCTB and emphasize the potential role of HLA-DR as a promising biomarker for TB diagnosis.

## 1. Introduction

Tuberculosis (TB) is the leading cause of death amongst human immunodeficiency virus (HIV-1)-infected individuals [1]. Moreover, 15 to 20% of all TB cases in developing countries are accounted for by extrapulmonary TB (EPTB) [2,3] which disproportionately affects immunocompromised patients [4,5]. Pericardial TB (PCTB), a severe form of EPTB, is the most common cause of pericarditis in TB endemic countries in Africa and Asia [6–8]. PCTB related morbidity is significant, with mortality (which generally occurs early in the onset of the disease), as high as 26% and increasing to approximately 40% in cohorts of predominantly HIV-infected persons [9,10].

HIV impairs both innate and adaptive immune responses, with the most obvious immune defect being a progressive reduction in absolute CD4+ T cell numbers and systemic hyper activation [11]. HIV-1 has also been shown to alter the balance of Mtb-specific T helper subsets, through the reduction of Th17 cells and T regulatory (Treg) cells [12–14], suggesting that HIV shifts Mtb-specific responses toward a more pathogenic/inflammatory profile [12]. Pulmonary TB-induced systemic inflammation has been studied extensively showing high concentrations of acute phase proteins and pro-inflammatory cytokines including C-reactive protein (CRP), serum amyloid P component (SAP), interferon gamma (IFN-γ), interferon gamma-induced protein 10 (IP-10), chemokine (C-C motif) ligand 1 (CCL1) and tumor necrosis factor alpha (TNF-α) in serum/plasma of active TB participants in comparison to other respiratory diseases, LTBI or healthy controls [15–18]. Furthermore, in patients with pulmonary TB admitted to intensive care units, serum levels of inflammatory factors such as interleukin (IL)-1, IL-6, IL-10, IL-12, and IL-4 are upregulated compared to healthy controls [19]. Based on these results several host inflammatory marker signatures have been proposed as biomarkers for TB diagnosis and the monitoring of treatment response, with superior performance compared to smear microscopy [15,16,20,21].

However, the influence of HIV-1 co-infection on the immune response to *Mtb* in the context of pulmonary and extrapulmonary TB remains poorly understood. Moreover, studies assessing immune responses at site of disease are scarce [22–24]. These studies reported higher levels of cytokines/chemokines at the site of disease in comparison to paired peripheral blood with exception of a few analytes, such as interferon gamma (IFN-γ), IL-1β and IL-8 which were reported to be significantly higher in peripheral blood instead [22–24]. Thus, in the current study, we measured 39 soluble markers in blood and at site of disease (pericardial fluid) to 1) compare the systemic cytokine environment between pulmonary and pericardial TB (PCTB) patients coinfected with HIV-1, 2) define the relationship between HIV viral load and the inflammatory profiles, 3) define whether peripheral inflammation signatures mirrors those at site of infection, 4) assess the impact of TB treatment on systemic inflammation and 5) evaluate the performance of previously described blood-based biomarkers to discriminate latent from active TB.

## 2. Materials and methods

### 2.1. Study population

Participants included in this study (n = 54) were recruited from the Ubuntu Clinic, Site B, Khayelitsha or the Groote Schuur Hospital Cardiology Unit (Cape Town, South Africa) between June 2017 and April 2019. Participants were divided in three groups according to their TB status: i) Pericardial tuberculosis (PCTB, n=18), ii) Pulmonary tuberculosis (PTB, n=20) and iii) Latent tuberculosis infection (LTBI, n=16).

The PCTB group (n = 18) included patients with either definite (Mtb culture positive in pericardial fluid (PCF), n = 9) or probable PCTB (n = 9). Probable PCTB was defined based on evidence of pericarditis with microbiologic confirmation of Mtb-infection elsewhere in the body and/or an exudative, lymphocyte predominant pericardial effusion with elevated adenosine deaminase (≥35 U/L), according to Mayosi et al [25]. Only three PCTB patients were HIV negative. Paired PCF and Blood were collected at the same time for PCTB patients. Patients from the PTB group (n = 20) were all HIV positive, tested sputum Xpert MTB/RIF (Xpert, Cepheid, Sunnyvale, CA) positive and had clinical symptoms and/or radiographic evidence of tuberculosis. All were infected by drug sensitive isolates of Mtb and had received no more than one dose of anti-tubercular treatment (ATT) at the time of baseline blood sampling. The LTBI group (n = 16) were all asymptomatic, had a positive IFN-γ release assay (IGRA, QuantiFERON-TB Gold In-Tube, Qiagen, Hilden, Germany), tested sputum Xpert MTB/RIF negative and exhibited no clinical evidence of active TB. All LTBI participants were HIV positive. Clinical characteristics of the study participants are shown in **Table 1**. Sputum and PCF Mtb culture, CD4 count, and HIV VL were performed by the South African National Health Laboratory Services. Active TB patients (PTB or PCTB) were followed up over the duration of their ATT and additional blood draws were performed at week 6 for PCTB, week 8 for PTB and week 24 for both diseased groups. All participants were adults (age ≥ 18 years) and provided written informed consent. The study was approved by the University of Cape Town Human Research Ethics Committee (050/2015 and 271/2019).

**Table 1.**
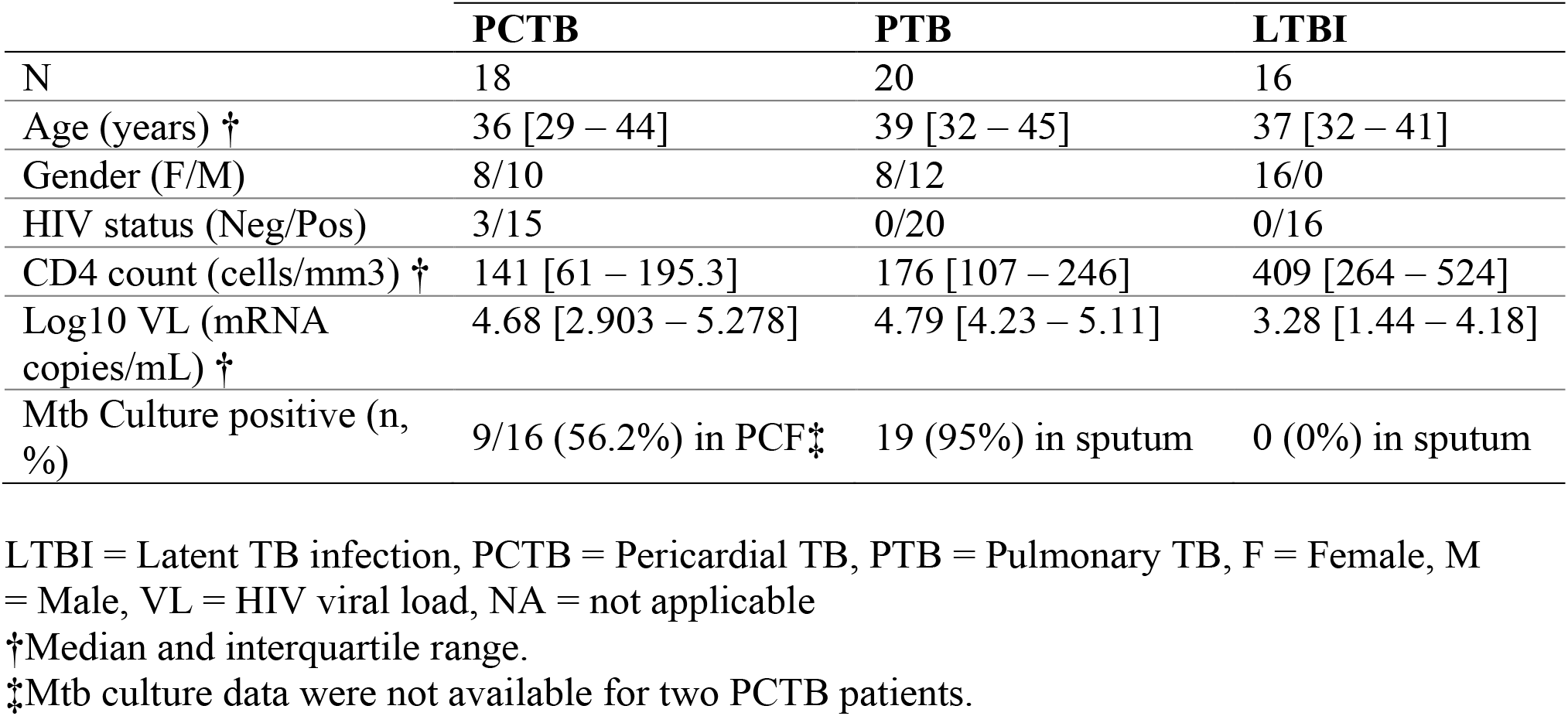
Clinical characteristics of study participants.

|  | **PCTB** | **PTB** | **LTBI** |
| --- | --- | --- | --- |
| N | 18 | 20 | 16 |
| Age (years) † | 36 [29 – 44] | 39 [32 – 45] | 37 [32 – 41] |
| Gender (F/M) | 8/10 | 8/12 | 16/0 |
| HIV status (Neg/Pos) | 3/15 | 0/20 | 0/16 |
| CD4 count (cells/mm3) † | 141 [61 – 195.3] | 176 [107 – 246] | 409 [264 – 524] |
| Log10 VL (mRNA copies/mL) † | 4.68 [2.903 – 5.278] | 4.79 [4.23 – 5.11] | 3.28 [1.44 – 4.18] |
| Mtb Culture positive (n, %) | 9/16 (56.2%) in PCF‡ | 19 (95%) in sputum | 0 (0%) in sputum |

LTBI = Latent TB infection, PCTB = Pericardial TB, PTB = Pulmonary TB, F = Female, M = Male, VL = HIV viral load, NA = not applicable
†Median and interquartile range.
‡Mtb culture data were not available for two PCTB patients.

### 2.2. Pericardial fluid, blood collection and whole blood assay

Pericardial fluid was obtained at the time of pericardiocentesis, placed in sterile Falcon tubes and transported to the laboratory at 4°C. Blood was collected in sodium heparin tubes and processed within 3 hours of collection. The whole blood or whole PCF assay were adapted from the protocol described by Hanekom et al [26]. Briefly, 0.5 mL of whole blood or 1 mL of whole PCF were stimulated with a pool of 300 Mtb-derived peptides (Mtb300, 2 μg mL^−1^) [27] at 37°C for 5 hours in the presence of the co-stimulatory antibodies, anti-CD28 and anti-CD49d (1 μg mL^−1^ each; BD Biosciences, San Jose, CA, USA) and Brefeldin-A (10 μg mL^−1^; Sigma-Aldrich, St Louis, MO, USA). Unstimulated cells were incubated with co-stimulatory antibodies and Brefeldin-A only. Red blood cells were then lysed in a 150 mM NH_4_Cl, 10 mM KHCO_3_ and 1 mM Na_4_EDTA solution. Cells were stained with a Live/Dead near-infrared dye (Invitrogen, Carlsbad, CA, USA) and then fixed using a transcription factor fixation buffer (eBioscience, San Diego, CA, USA), cryopreserved in freezing media (50% fetal calf serum, 40% RPMI and 10% dimethyl sulfoxide) and stored in liquid nitrogen until use.

### 2.3. Cell staining and flow cytometry

Cryopreserved cells were thawed, washed and permeabilized with a transcription factor perm/wash buffer (eBioscience). Cells were then stained at room temperature for 45 min with the following antibodies: CD3 BV650 (OKT3; BioLegend, San Diego, CA, USA), CD4 BV785 (OKT4; BioLegend), CD8 BV510 (RPA-T8; BioLegend), HLA-DR BV605 (L243; BioLegend), IFN-γ BV711 (4S.B3; BioLegend), TNF-α PE-Cy7 (Mab11; BioLegend eBioscience) and IL-2 PE/Dazzle (MQ1-17H12; BioLegend). Samples were acquired on a BD LSR-II and analysed using FlowJo (v10.8.1, FlowJo LCC, Ashland, OR, USA). A positive cytokine response was defined as at least twice the background of unstimulated cells. To define the phenotype of Mtb300-specific CD4 T cells, a cut-off of 30 events was used.

### 2.4. Luminex® Multiplex Immunoassay

Using Luminex® technology, we measured the levels of 39 analytes using antibodies supplied by Merck Millipore (Billerica, Massachusetts, USA) and R&D Systems (Minneapolis, MN, USA). The analytes measured included: Granzyme B (GrB), interleukin 2 (IL-2), interleukin 8 (IL-8), interleukin 12p40 (IL-12p40), macrophage colony-stimulating factor (M-CSF), tumor necrosis factor alpha (TNF-α), transforming growth factor beta (TGF-β), complement component 3 (C3), complement component 4 (C4), C-reactive protein (CRP), serum amyloid P (SAP), interleukin 22 (IL-22), Galectin-3 (Gal-3), intercellular adhesion molecule 1 (ICAM-1), neural cell adhesion molecule 1 (NCAM-1), granulocyte colony-stimulating factor (G-CSF), interferon gamma (IFN-γ), interleukin 6 (IL-6), interleukin 10 (IL-10), interleukin 27 (IL-27) and vascular endothelial growth factor (VEGF), monokine induced by gamma (MIG), monocyte chemoattractant protein 2 (MCP-2), granulocyte chemoattractant protein 2 (GCP-2), chemokine (C-X-C motif) ligand 11 (CXCL11), macrophage inflammatory protein 1 beta (MIP-1β), chemokine (C-C motif) ligand 1 (CCL1) and interferon gamma-induced protein 10 (IP-10), cluster of differentiation 163 (CD163), interleukin 6 receptor alpha (IL-6Rα), cluster of differentiation 30 (CD30), interleukin 2 receptor alpha (IL-2Rα), apolipoprotein A-I (ApoA-I), apolipoprotein C-III (Apo-CIII), oncostatin M (OSM), interleukin 33 receptor (IL-33R), osteopontin (OPN), platelet derived growth factor BB (PDGF-BB) and thrombomodulin (TM). All samples were evaluated undiluted or diluted according to the manufacturer’s recommendations. Samples were randomized to assay plates with the experimenter blinded to sample data. All assays were performed and read at UCT on the Bio-Plex platform (Bio-Rad), with the Bio-Plex Manager Software (v6·1) used for bead acquisition and analysis.

### 2.5. Statistical Analyses

Statistical tests were performed in Prism (v9.1.3, GraphPad Software Inc, San Diego, CA, USA). Non-parametric tests were used for all comparisons. The Kruskal-Wallis test with Dunn’s multiple comparison test was used for multiple comparisons, the Spearman rank test for correlation and the Mann-Whitney and Wilcoxon matched pairs test for unmatched and paired samples, respectively. When the measured analyte was below the limit of detection in more than 20% of the samples (i.e., M-CSF and IL-10), the analyte was not included in the correlation with plasma HIV VL and HLA-DR expression on Mtb-specific CD4 T cells. Unsupervised hierarchical clustering analysis (HCA, Ward method), principal component analyses (PCA) were carried out in JMP (v16.0.0; SAS Institute, Cary, NC, USA). For HCA and PCA, the min-max normalization method (i.e., feature scaling, analyte value - min / max - min) was used to scale data in the 0 to 1 range. The predictive abilities of combinations of analytes were investigated by general discriminant analysis (GDA) in JMP. The diagnostic ability of HLA-DR expression on Mtb-specific CD4 T cells were assessed by receiver operator characteristics (ROC) curve analysis. Optimal cut off values and associated sensitivity and specificity were determined based on the Youden’s Index [28]. Analyte network analysis was performed using Gephi (v0.9.2, University of Technology of Compiègne, Compiègne, France). The Bonferroni method [29] was used to adjust for multiple comparisons. A p-value of <0.05 was considered statistically significant.

## 3. Results

### 3.1 Study population

The clinical characteristics of participants are presented in **Table 1**. Participants (n = 54) were classified into three groups according to their TB status: PCTB (n = 18), PTB (n = 20) and LTBI (n = 16). Median age was comparable between the three groups. All participants were HIV-infected except for three PCTB patients. LTBI participants had a lower plasma HIV-1 viral load (VL) and higher absolute CD4 count compared to the PCTB and PTB groups (median Log_10_ VL: 3.28 vs 4.68 and 4.79 copies mL^−1^, respectively and median CD4: 409 vs 141 and 176 cells mm^−3^, respectively, **Table 1**).

### 3.2 Comparison of the systemic inflammatory profile between LTBI, PTB and PCTB

Plasma levels of 39 analytes, including cytokines, chemokines, apolipoproteins, chemokine, protein receptors, and fibrosis-related analytes, were measured in all participants (the complete list of measured analytes is presented in the material and methods section). Assessing the overall systemic inflammatory profile using unsupervised hierarchical clustering (**Fig. 1a**) and principal component analysis (**Fig. 1b**) we showed an evident separation between LTBI and active TB participants (PCTB and PTB), driven by elevated levels of most of the measured inflammatory markers. However, there was no noticeable separation between the PCTB and PTB groups, suggesting comparable systemic inflammation in these groups. Individual analysis of measured analytes showed that 15 markers were significantly higher in both PTB and PCTB compared to the LTBI group, including innate-related inflammation markers (such as IL-6, TNF-α, and IL-8), acute phase protein (CRP) and chemokines (CCL1, MIG, IP-10 and CXCL11). VEGF also showed a similar profile, with the p-value between LTBI and PTB being borderline significant (p = 0.0503) (**Supplementary fig. 1 and Supplementary table 1**). IL-6Rα and G-CSF were the only markers that were observed to be differentially expressed between PTB and PCTB (**Supplementary fig. 1 and Supplementary table 1**), highlighting similarities between the different clinical forms of TB. Only one marker, OPN showed increased expression levels only in the PCTB group compared to LTBI (p = 0.0063) while no significant difference was observed for the PTB group (p = 0.374) (**Supplementary fig. 1 and Supplementary table 1**). Elevated OPN levels have been associated with severe tuberculosis [30]. Next, we defined the interplay between markers, using network analysis (Fruchterman-Reingold algorithm, **Fig. 1c**). In LTBI participants, TNF-α and MIP-1β were the most central nodes, showing the most connections (positive associations) with other analytes. In active TB patients (both PTB and PCTB), the network structure was substantially altered; and while MIP-1β remained a predominant node, TGF-β emerged as a new influential node, with multiple negative associations with analytes such as IL-12p40, ApoA-I or G-CSF (**Fig. 1c**). Overall, these results illustrate that active TB disease significantly increases systemic inflammation and PCTB and PTB participants share similar inflammatory signatures.

**Figure 1.**
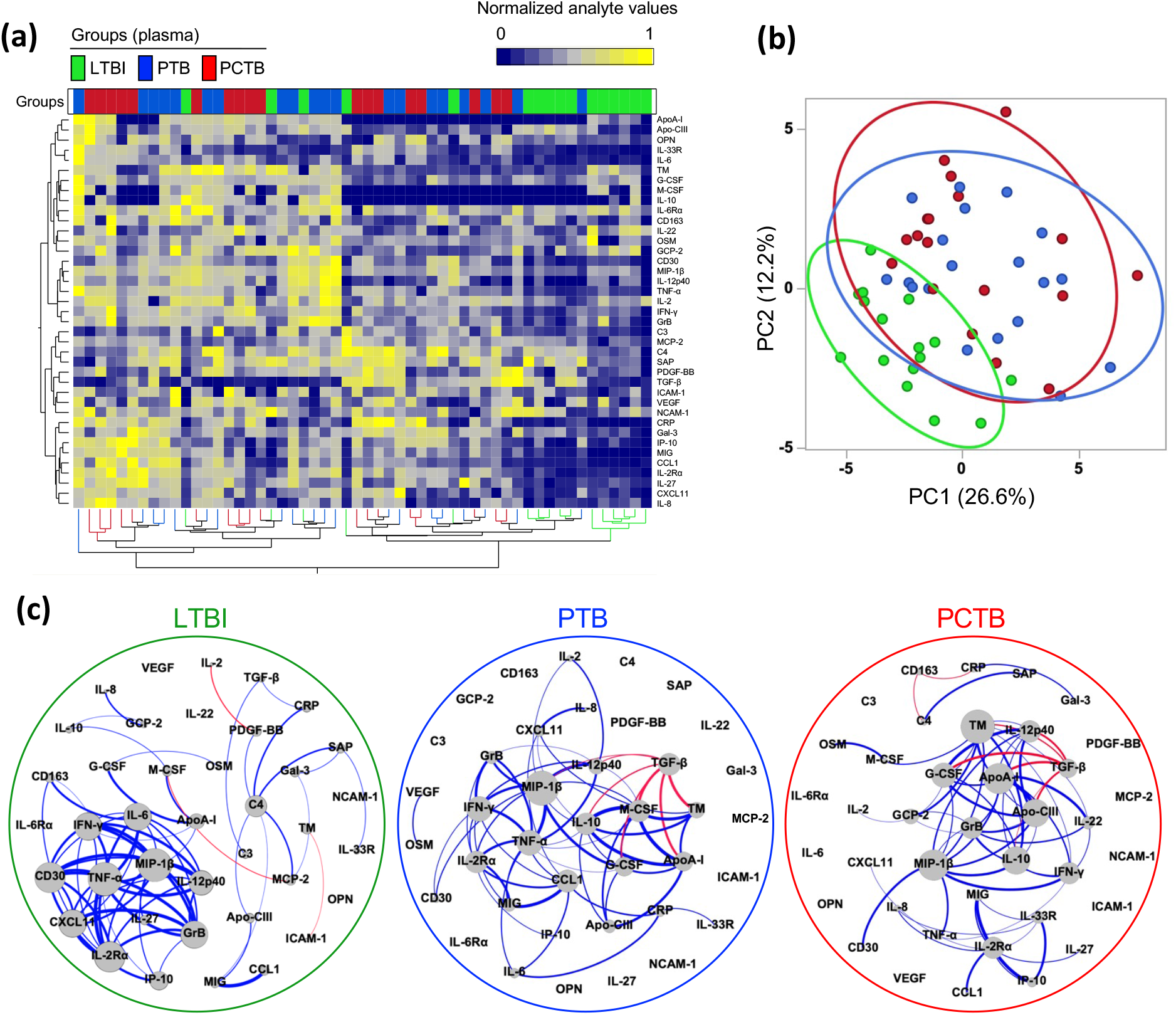
Analyte profiles in the different TB groups at baseline. **(a)** A non-supervised two-way hierarchical cluster analysis (HCA, Ward method) was employed to evaluate the TB groups using the 39 measured analytes. TB status (PCTB in red, PTB in blue and LTBI in green) of each patient is indicated at the top of the dendrogram. Data are depicted as a heatmap colored from minimum to maximum normalized values for each marker. **(b)** Principal component analysis (PCA) on correlations based on the 39 analytes was used to explain the variance of the data distribution in the cohort. Each dot represents a participant. The two axes represent principal components 1 (PC1) and 2 (PC2). Their contribution to the total data variance is shown as a percentage. **(c)** Analyte network analysis (Fruchterman-Reingold algorithm) in plasma of LTBI, PTB and PCTB participants. Size of nodes indicate the number of connections. Size of edges indicate the spearman r value (only r > 0.6 were included). Blue lines: positive correlation. Red lines: negative correlation.

### 3.3 Relationship between inflammatory profile and HIV viral load

To examine the interplay between HIV viral load (VL) and cytokine profile, we defined the associations between cytokine concentrations and HIV VL in plasma. Of the 39 measured analytes, 12 markers positively associated with HIV VL in the LTBI group (**Fig. 2a**). Several of those have been previously reported as HIV-associated systemic inflammation markers, including IL-2Rα [31], CXCL11 [32], IL-6 [33], IFN-γ [34], IP-10 [35], TNF-α [35], and CD30 [36]. In both the PTB and PCTB groups, most of these correlations were disrupted with six analytes correlating with HIV VL in the PTB group and only one in the PCTB group (**Fig. 2a**). The only cytokine which maintained significant correlation with HIV VL in all groups was IL-12p40, albeit the correlation strength was weaker in the diseased groups (r = 0.83, p = 0.0002 vs r = 0.49, p = 0.028 in the PTB group and r = 0.63, p = 0.012 in the PCTB group) (**Fig. 2b**). IP-10 concentration only showed a significantly positive correlation with HIV VL in the LTBI group (r = 0.82, p = 0.0002), and was largely disrupted in both the PTB and PCTB groups (r = 0.29, p = 0.26 and r = 0.25, p = 0.37, respectively) (**Fig. 2b**). No negative associations were observed in the LTBI and PTB groups, however, TGF-β showed a strong negative association with HIV VL in the PCTB group (r = −0.65, p = 0.0133) (**Fig. 2a**). These findings suggest that active TB disease disrupts HIV-associated systemic inflammation.

**Figure 2.**
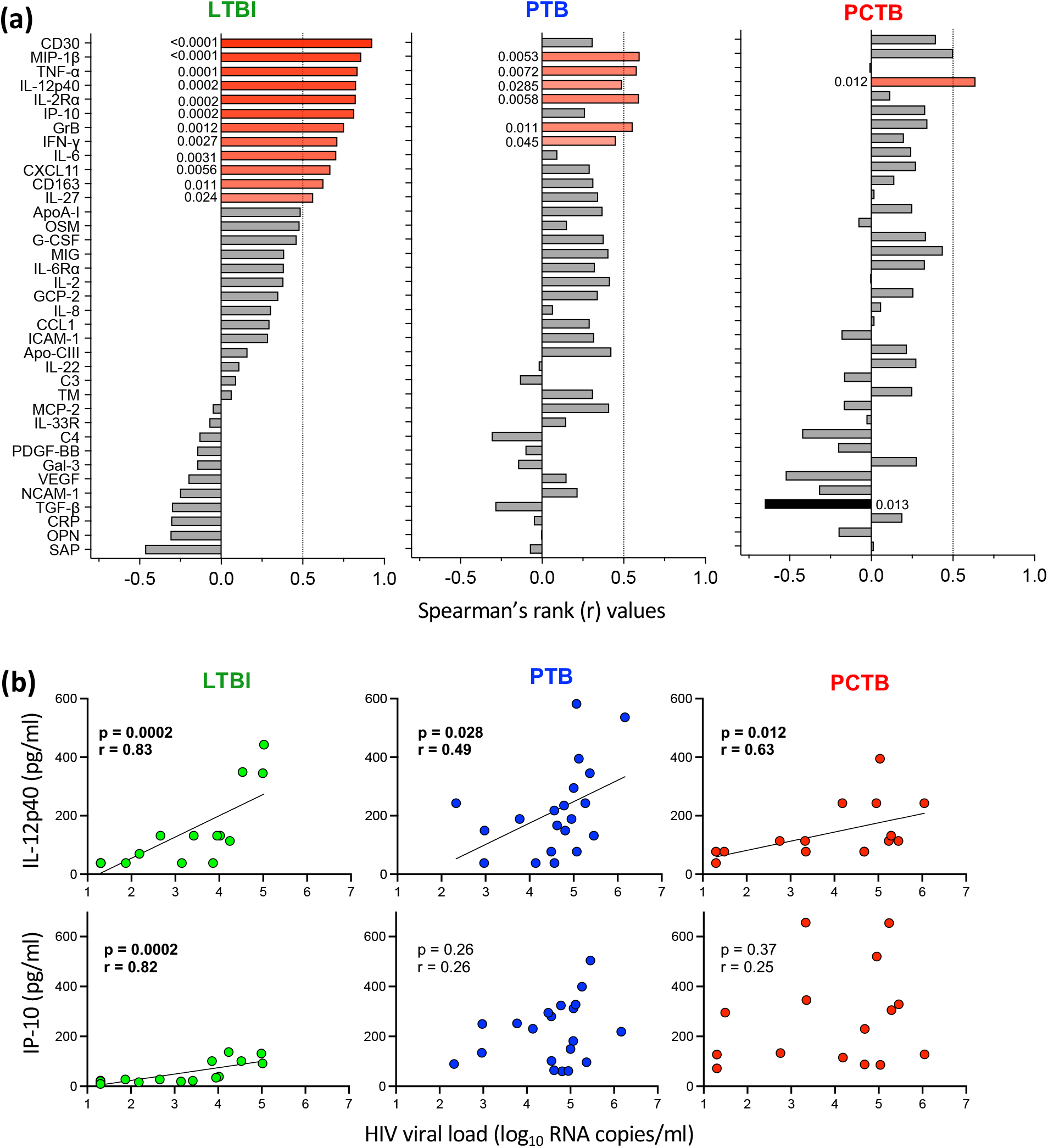
Univariate associations between HIV VL and analyte concentrations in the different TB groups. **(a)** Spearman’s rank values of the univariate correlation between each analyte and the HIV VL in LTBI participants, PTB participants, and PCTB participants plasma samples. Red bars indicate positive correlations, Black bars indicate negative correlations, and grey bars indicate non-significant correlations. **(b)** Depicts the examples of IL-12p40 (maintained relationship between the TB groups) and IP-10 (disrupted relationship between the TB groups). The line indicates linear regression for statistically significant correlations.

### 3.4 Profile of soluble markers in plasma compared to pericardial fluid

To better understand compartmentalization, we compared the profiles of expression of the 39 measured analytes in plasma and PCF from PCTB participants, using hierarchical clustering analysis and PCA (**Fig. 3a and b**). There was a clear separation between sample types, where PC1 accounted for 42% and PC2 11.2% of the variance (**Fig. 3b**). Furthermore, visualizing sample clustering using a constellation plot, we observed that cluster 2 (comprised of PCF samples) was divided into 2 distinct sub-clusters, where cluster 2b was enriched in participants who were PCF culture positive (5/7, 72%) compared to patients included in cluster 2a (4/12, 33%) (**Fig. 3c**). However, looking at individual analytes, we did not find significant difference between PCF culture negative and PCF culture positive samples (data not shown).

**Figure 3.**
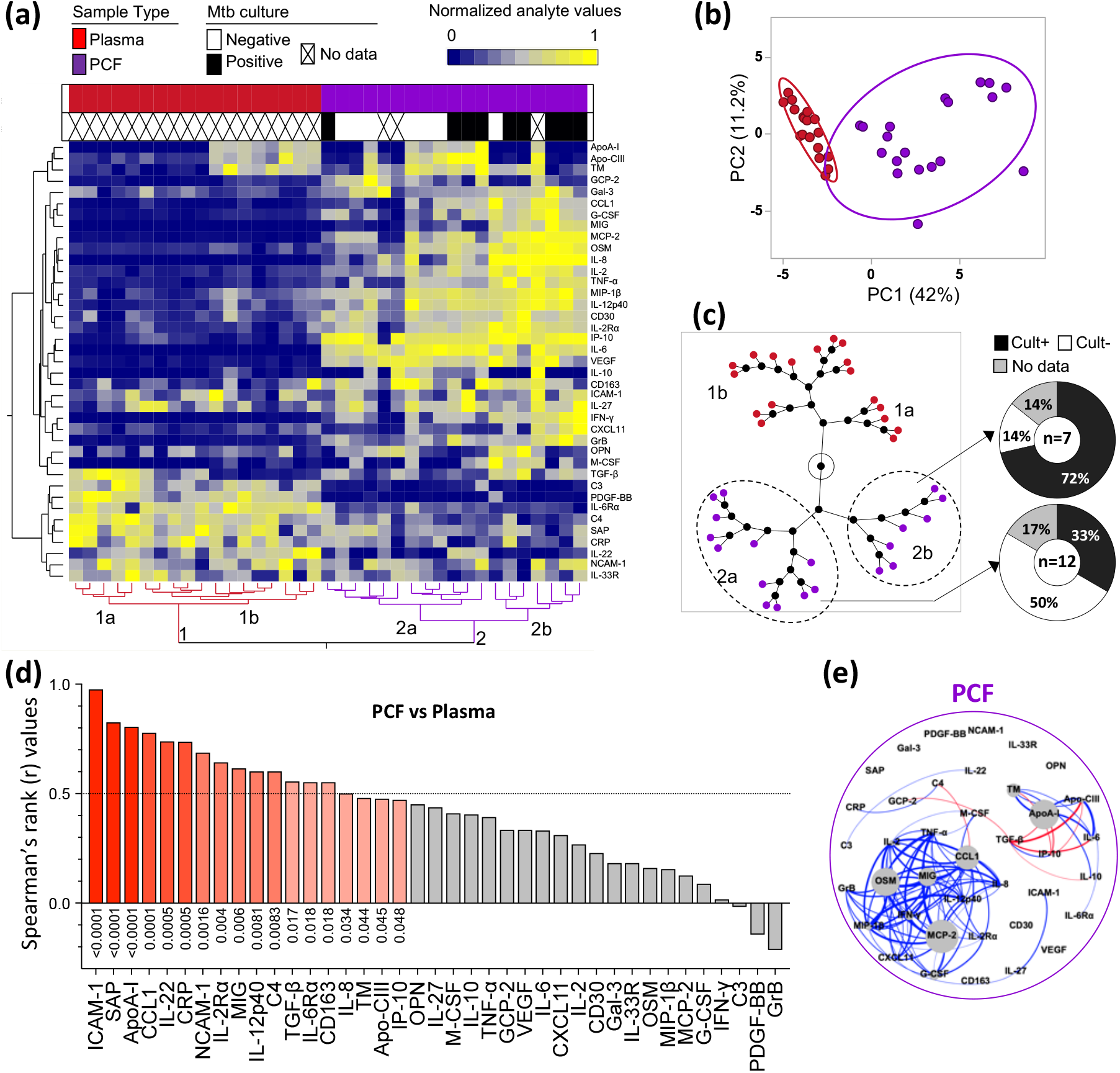
Analyte profiles in peripheral blood (Plasma) and site of disease (Pericardial fluid) in PCTB participants. **(a)** A non-supervised two-way hierarchical cluster analysis (HCA, Ward method) was employed to evaluate the two sites using the 39 analytes. The sample type and Mtb culture results (PCF in purple, Plasma in red; Mtb culture negative in white and positive in black) of each patient is indicated at the top of the dendrogram. Data are depicted as a heatmap colored from minimum to maximum normalized values detected for each marker. **(b)** Principal component analysis (PCA) on correlations based on the 39 analytes was used to explain the variance of the data distribution in the subgroup. Each dot represents a participant. The two axes represent principal components 1 (PC1) and 2 (PC2). Their contribution to the total data variance is shown as a percentage. **(c)** Constellation Plot-cluster analysis based on all measured analytes. Each dot represents a participant and is color-coded according to sample type. Each cluster obtained for the HCA is identified by a number. **(d)** Pairwise correlation of the 39 analytes. Red bars indicate a positive correlation, Black bars indicate a negative correlation, and grey bars indicate a non-significant correlation. **(e)** Analyte network analysis in PCF of PCTB participants. Size of nodes indicate the number of connections. Size of edges indicate the spearman r (only r > 0.6 were included). Blue lines: positive correlation. Red lines: negative correlation.

Univariate analysis of analytes showed that the concentrations of 25 out of the 39 measured analytes were significantly higher in PCF in comparison to paired plasma samples, only 9/39 were significantly higher in plasma compared to PCF, and 5/39 showed no significant difference in expression between the two sample types after correction of the p-values for multiple testing (**Supplementary fig. 2 and Supplementary table 2**).

To better understand the relationship between peripheral and site of disease inflammation, pairwise comparisons (plasma vs PCF) were assessed. Significant positive correlations were observed for 18 out of the 39 analytes (with r and p ranging from 0.98 - 0.47 and <0.0001 - 0.048, respectively), the highest Spearman’s rank r values for significant positive correlations were observed for ICAM-1, SAP, and ApoA-I (**Fig. 3d**). A summarized representation of the associations between plasma and PCF for each analyte is shown in **fig. 3d** and individual correlation plots of all the significant associations are presented in **supplementary fig. 3**. We then defined the interplay between markers in PCF, using network analysis (Fruchterman-Reingold algorithm, **Fig. 3e**). OSM, MCP-2 and ApoA-I were the most central nodes, with OSM and MCP-2 showing positive associations with other analytes. While ApoA-I showing mostly negative associations with analytes such as TGF-β, IP-10 and Apo-CIII (**Fig. 3e**). Overall, these results show that inflammatory response at site of disease was greater than in blood. However, inflammatory profile in PCF partially mirrored inflammatory events in blood.

### 3.5 Associations between systemic inflammation and the activation of Mtb-specific CD4+ T cells in blood and at site of disease

HLA-DR expression on peripheral Mtb-specific CD4+ T cells has been shown to discriminate latent from active TB infection [37–39]. To better understand the relationship between inflammation and T cell activation, we measured the expression of HLA-DR on Mtb-specific CD4+ T cells in blood from LTBI, PTB, PCTB and PCF from PCTB participants. As expected, HLA-DR expression on peripheral Mtb-specific CD4+ T cells was significantly higher in the aTB groups (PTB and PCTB) compared to LTBI (medians: 62.30% and 70.85% vs 17.20%, respectively, p >0.0001). Moreover, HLA-DR expression on Mtb-specific CD4+ T cells in PCF was significantly higher compared to blood in the PCTB group (medians: 78.30% vs 69.90%, respectively, p= 0.0341) (**Fig. 4a and b**). We then assessed the association of HLA-DR expression on Mtb-specific CD4 T cells and the concentrations of each measured analyte at the site of disease (PCF) and in blood from PCTB participants as well as blood from PTB participants (**Fig. 4c**). At disease site, we observed positive associations between HLA-DR expression on Mtb-specific CD4 T cells and 10 analytes, including CCL1, G-CSF, OSM, IL-8, IL-2 and IL-2Rα (with r value > than 0.6).

**Figure 4.**
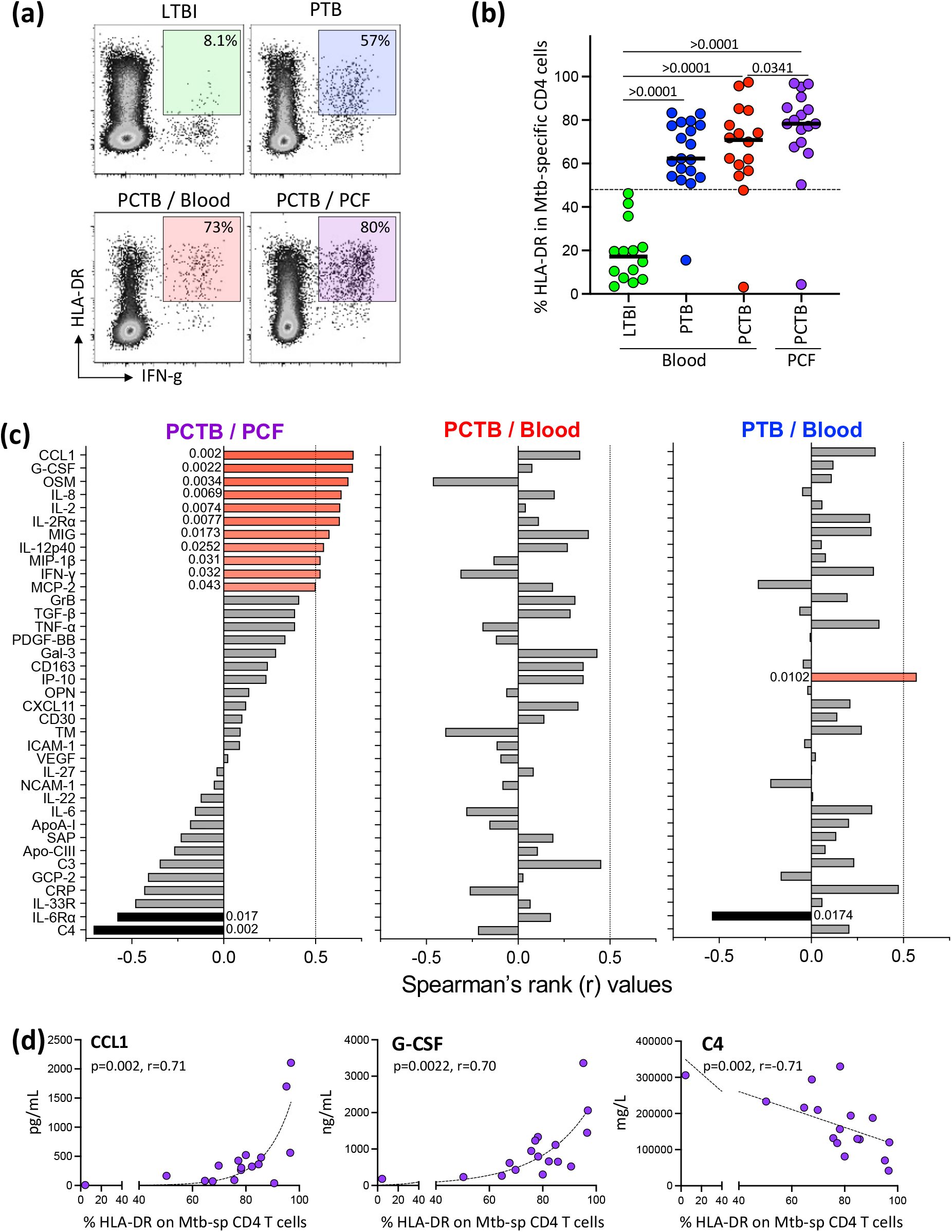
Univariate associations between HLA-DR and analyte concentrations in the different TB groups. **(a)** Representative flow cytometry plots of the expression of HLA-DR. **(b)** Expression of HLA-DR on Mtb-specific CD4 T cells in response to Mtb300. **(c)** Spearman’s rank values of the univariate correlation between each analyte and between Mtb-specific CD4 T cell activation (HLA-DR) level at the site of disease (PCF) in PCTB participants, in blood of PCTB and PTB participants, respectively. Red bars indicate a positive correlation, Black bars indicate a negative correlation, and the grey bars indicate non-significant correlation. **(d)** Representative graphs showing the positive (CCL1 and G-CSF) and negative (C4) correlation to HLA-DR frequency at the site of disease (PCF). Statistical comparisons were performed using a Kruskal-Wallis test, adjusted for multiple comparisons (Dunn’s test) for blood LTBI vs PTB vs PCTB, Wilcoxon test for blood PCTB vs PCF PCTB and the Mann-Whitney test to compare blood LTBI and PCF PCTB.

Negative associations were observed with C4 (r = −0.71, p = 0.002) and IL-6Rα (r = −0.54, p = 0.017) (**Fig. 4d**). None of these associations were observed in peripheral blood (**Fig. 4c**). In PTB participants, HLA-DR expression on peripheral Mtb-specific CD4+ T cells associated with only 2 analytes, namely IP-10 (r = 0.57, p = 0.0102) and IL-6Rα (r = −0.54, p = 0.0174) (**Fig. 4c**). These data suggest a coordinated and compartmentalized immune response at the disease site.

### 3.6 Impact of TB treatment on the inflammatory profile in plasma

Monitoring of TB treatment response is challenging mainly due to the lack of specific and sensitive blood-based tools. In the current study, we examined the effect of TB treatment on the expression of inflammation markers. First, we compared the overall systemic inflammatory profile in participants with LTBI and in aTB patients (PTB and PCTB) 24 weeks after TB treatment initiation using unsupervised hierarchical clustering (**Fig. 5a**) and principal component analysis (**Fig. 5b**). No specific clustering was observed between the groups, showing a global normalization of the inflammation signature at treatment completion. Furthermore, we performed univariate analysis comparing the level of expression of each analyte at baseline (before TB treatment initiation), week 6 or 8 and week 24 post treatment initiation (**Supplementary fig. 4 and Supplementary table 3**). Of the 39 measured analytes, 13 showed significant reduction between baseline, week 6/8 and/or week 24 in both the PTB and PCTB groups (**Supplementary fig. 4a and Supplementary table 3**). An additional eight analytes showed reduction between the three time points in the PTB group only (**Supplementary fig. 4b and Supplementary table 3**).

**Figure 5.**
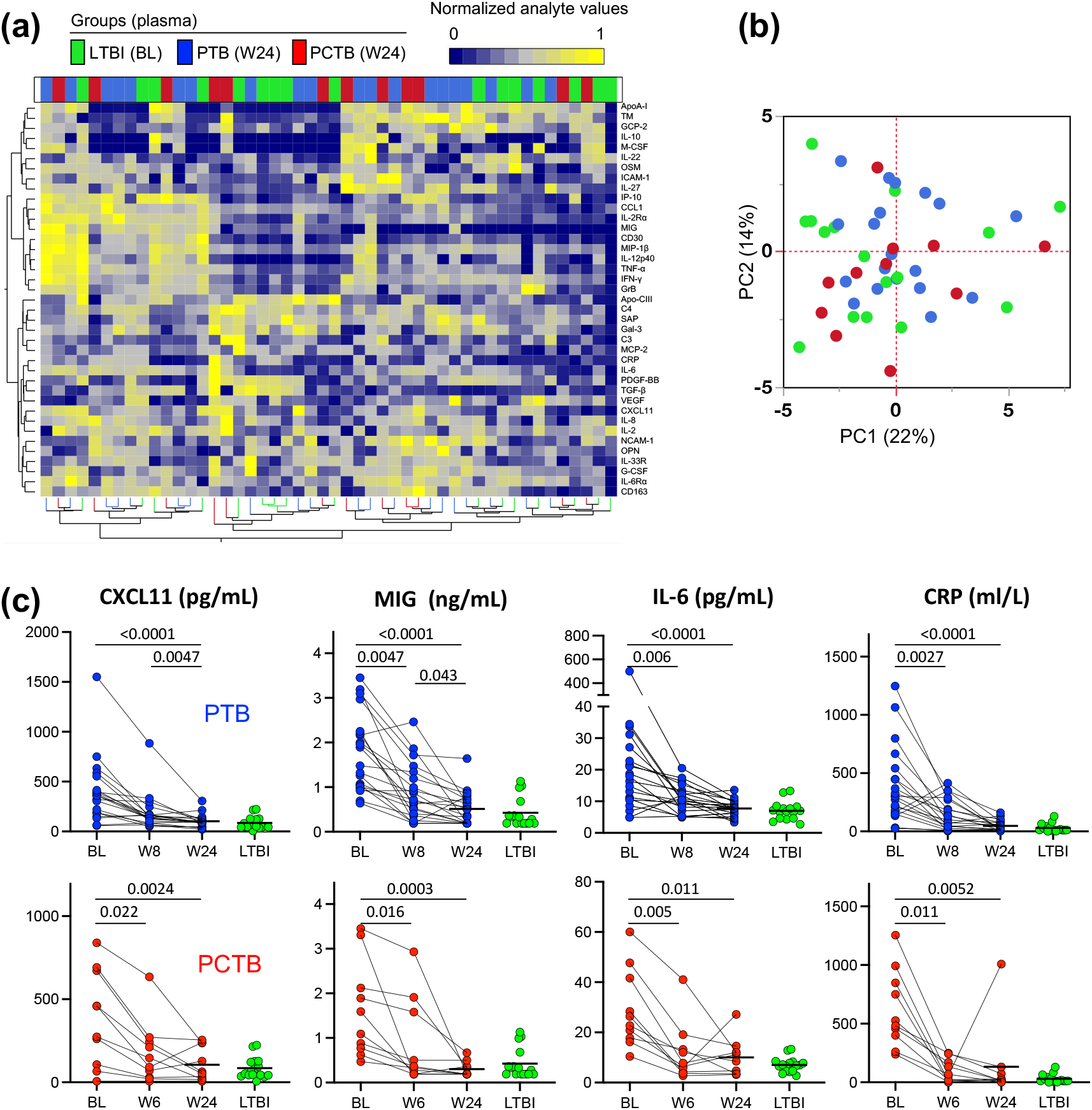
Analyte profiles in the different TB groups before, during and post TB treatment. **(a)** A non-supervised two-way hierarchical cluster analysis (HCA, Ward method) was employed to grade the TB groups using the 39 analytes. TB status (PCTB in red, PTB in blue and LTBI in green) of each patient is indicated at the top of the dendrogram. Data are depicted as a heatmap colored from minimum to maximum normalized values detected for each marker. **(b)** Principal component analysis (PCA) on correlations based on the 39 analytes was used to explain the variance of the data distribution in the cohort. Each dot represents a participant. The two axes represent principal components 1 (PC1) and 2 (PC2). Their contribution to the total data variance is shown as a percentage. **(c)** Representative graphs showing the change of concentrations of CXCL11, MIG, IL-6 and CRP with treatment and no statistical difference between week 24 post-treatment initiation and LTBI in both PTB and PCTB groups, respectively. Statistical comparisons were performed using a Friedman test, adjusted for multiple comparisons (Dunn’s test) for BL v W6/W8, BL v W24 and W6/W8 v W24 and the Mann-Whitney test to compare LTBI with W24, p-values were adjusted using the Bonferroni method.

Representative plots of analytes including, CXCL11, MIG, IL-6 and CRP depict the significant reduction of expression of analytes with TB treatment from baseline, week 6/8 to end of treatment (week 24) in both PTB and PCTB groups (**Fig. 5c**). These data suggest that the overall inflammatory profile normalized upon TB treatment completion in both PTB and PCTB.

### 3.7 Comparison of HLA-DR expression and biosignatures derived from soluble analytes in discriminating LTBI from active TB

Previous studies have shown the potential of blood-based markers to distinguish LTBI from aTB, including biosignatures derived from soluble markers and HLA-DR expression on MTB-specific T cells [15,16,20,21,37,38]. Although this study was not designed as a diagnostic study, we explored this aspect, wherein we assessed the ability of HLA-DR expression to distinguish LTBI from PTB, PCTB or any aTB (PTB + PCTB) and compared it with previously described biosignatures that included analytes measured in this study. We generated receiver operating characteristic (ROC) curves from data obtained in Mtb-specific CD4 T cells. Consistent with previous reports, HLA-DR expression on Mtb-specific CD4 T cells showed a great capability to distinguish LTBI from PTB (p<0.0001, area-under-the-curve (AUC) = 0.97, 95% CI: 0.92 – 1.00, sensitivity: 97.75%, specificity: 100%, at an optimal cut-off of 48.5%) (**Supplementary fig. 5a and b**). Moreover, HLA-DR expression also discriminated LTBI from PCTB (p<0.0001, AUC = 0.94, 95% CI: 0.82 – 1.00, sensitivity: 93.75%, specificity: 100%, at an optimal cut-off of 46.9%) and LTBI from any aTB (p<0.0001, AUC = 0.96, 95% CI: 0.90 – 1.00, sensitivity: 94.29%, specificity: 100%, at an optimal cut-off of 46.9%) (**Supplementary fig. 5a and b**).

We assessed the performance of previously described soluble biosignatures our data set to and compared soluble biosignature performance to HLA-DR expression. We identified six different published biosignatures which include analytes measured in this study: [IL-12p40 + IL-10] [21], [IFN-γ + IL-10 + IL-12p40] [21], [TNF-α + IL-12p40] [21], [CCL1 + CRP] [15], [CCL1 + TNF-α] [16], and [IL-6Rα + IL-2Rα] [20].

These biosignatures discriminated LTBI from PTB with AUCs ranging from 0.72-0.9 and corresponding sensitivity and specificity ranging from 55% − 85% and 75% - 100%, respectively. They also discriminated LTBI from PCTB with AUCs ranging from 0.64 - 1.00 and corresponding sensitivity and specificity ranging from 61.11% - 83.33% and 62.5% - 93.75%, respectively, while they discriminated LTBI from any aTB (PTB + PCTB) with AUCs ranging from 0.69 - 0.98 and corresponding sensitivity and specificity ranging from 52.63% - 76.32% and 62.50% - 100%, respectively (**Supplementary table 4**). Detailed performances of these signatures in comparison to HLA-DR expression are shown in **supplementary table A.4**.

None of these biosignatures out-performed HLA-DR expression in discriminating LTBI from the diseased groups (**Supplementary table 4**). These findings suggest that HLA-DR is a better biomarker than soluble markers for discriminating between the different TB groups.

## 4. Discussion

EPTB represents a small but significant proportion of all TB cases globally, particularly in HIV-infected patients and is frequently difficult to diagnose. However, immune and inflammatory responses at the site of disease remains understudied. In this study, we compared the TB-associated inflammatory response in HIV-infected participants between latent, pulmonary, and pericardial TB infection. We also compared the inflammatory signature in blood and at site of disease (i.e., PCF) in PCTB patients. Moreover, we measured HLA-DR expression on Mtb-specific CD4 T cells from whole blood and compared its diagnostic potential to previously described biosignatures derived from different combinations of soluble markers.

We show that PTB in HIV-infected patients is characterized by increased systemic inflammation compared to LTBI persons. This is in accordance with previous reports showing elevated inflammatory markers (such as CRP, IP-10, IFN-γ, CCL1, and VEGF) in unstimulated plasma or serum in aTB compared to LTBI or other respiratory diseases regardless of HIV status [15,16,18]. In HIV negative individuals, distinct inflammatory profiles in PTB versus extra pulmonary TB have been reported, which were speculated to be the consequence of differences between disseminated versus more localized infection [40]. However, here, we observed a similar inflammatory profile in HIV-infected PTB individuals and HIV-infected PCTB individuals. These differences may be explained by the different analytes measured in the Vinhaes et al [40] study and the current study, with only seven analytes overlapping between the two studies (namely, IL-2, IL-6, IL-8, IL-10, IL-27, TNF-α, and IFN-γ). Moreover, the Vinhaes et al [40] study included patients with different types of EPTB (including Pleural TB, TB lymphadenitis and Miliary TB) while our study focused exclusively on PCTB patients.

To improve our understanding of immunological mechanisms at the disease site, we compared inflammatory profile at disease site and in plasma. A study by Matthews et al [22], assessing the inflammatory response at the disease site, showed compartmentalization of inflammatory proteins (including IL-6, IL-8 and IFN-γ) in PCF compared to blood. Our results are in accordance with this study, showing that inflammation was greater at the site of disease compared to the periphery and further demonstrate that there was a partial mirroring of the innate-associated inflammatory response (such as CCL1, IL-12p40, TGF-β and IL-8) between blood and disease site. Interestingly, Th1 cytokines levels (IFN-γ and IL-2) in PCF did not correlate with plasma levels. We previously reported that there was no correlation between the frequency of Mtb-specific CD4 T cells in blood and PCF [41] and recent data from murine model suggests that the rate of migration of T cell to the disease site is mostly regulated by the pattern of chemokine receptors they expressed [42].

TB diagnosis is challenging due to the lack of rapid, accurate, blood-based diagnostic tests. HLA-DR expression on Mtb-specific CD4 T cells has been shown to be a robust marker in discriminating latent TB from aTB [37–39] and EPTB [43]. In this study, we observed HLA-DR to be significantly highly expressed in blood of aTB compared to LTBI, it was also highly expressed, at the site of disease (PCF) in PCTB participants compared to blood of the same participants. Our findings are in agreement with previously published studies [37–39,43] and further suggest that the extent of activation of infiltrating CD4 T cells associate with the inflammatory profile at the disease site.

Several biosignatures consisting of host soluble inflammatory markers have been described as promising tools for TB diagnosis [15,16,20,21]. Here, we used our cohort as a validation cohort to compare their performance in discriminating LTBI from aTB, and several previously identified biosignatures continued to show promise in our cohort. However, none of these biosignatures showed better performance compared to the measure of HLA-DR expression on Mtb-specific CD4 T cells, which met the WHO target product profile (TPP) recommendations for a point of care non-sputum-based triage test [44]. These data further emphasize the role of HLA-DR as a promising biomarker for TB diagnosis.

Sputum culture conversion at two months post treatment initiation remains the most widely used tool for the evaluation of TB treatment response [45,46]. However, in individuals with PCTB who are sputum smear or culture negative for Mtb, monitoring of treatment response is solely assessed clinically as there are no validated blood biomarkers to assist in this regard. Changes in blood biomarker levels during antitubercular treatment in either PTB or EPTB cases has been previously reported in a number of prospective studies [18,47–58], showing the normalization of several inflammatory markers (such as CRP, IP-10, CCL1, IFN-γ and TNF-α) after successful TB treatment. Our findings are in accordance with these results and add to the current knowledge, showing that the concentrations of several of the biomarkers tested (21 out of 39 and 13 out of 39) decreased at treatment completion to levels observed in LTBI participants in both the PTB and PCTB groups, respectively. The discrepancy in the normalization of inflammatory profile after treatment between PTB and PCTB could be related to disease severity, where disseminated disease has been shown to present with elevated systemic bacterial burden and higher mortality [59] and limited drug penetration at the site of disease. Thus, our study confirms that measuring blood biomarkers may have utility to monitor treatment response in both pulmonary and extra-pulmonary TB. Our study has several limitations. First, most of the participants were HIV infected, we were thus unable to define the impact of HIV infection on TB-induced inflammatory profiles. Second, we did not have long-term follow-up clinical data to identify potential TB relapse, so long-term outcome could not be related to inflammatory profiles. Third, the current study was not designed to identify novel diagnostic markers, thus we confined our analysis to previously described blood-based biomarkers. However, further assessments of HLA-DR expression on Mtb-specific CD4 T cells are required in well-designed diagnostic studies. Finally, further experiments including patients with non-tuberculous pericardial effusion will be necessary to define whether the observed inflammatory signatures in plasma and at site of disease are TB specific. Regardless of the limitations, our results show that in a largely HIV-infected cohort with advanced immunosuppression, PCTB and PTB share similar inflammatory signature and aTB disrupts the relationship between HIV VL and soluble analytes. These results also reveal that profiles of markers at the site of disease are distinct from peripheral blood though some markers strongly correlate. Furthermore, upon completion of TB treatment, levels of soluble analytes normalized and lastly, we showed that in HIV-infected patients, assessing the expression of HLA-DR on Mtb-specific CD4 T cells had a better potential to discriminate PCTB and PTB from LTBI compared to biosignatures derived from soluble markers.

## Acknowledgments

The authors thank the study participants, the clinical staff at the Khayelitsha Site B Community Health Centre in Cape Town and the laboratory staff at the Wellcome Centre for Infectious Disease Research in Africa at the University of Cape Town.

## Funding

This work was supported by the European and Developing Countries Clinical Trials Partnership EDCTP2 programme; the European Union (EU)’s Horizon 2020 programme (Training and Mobility Action TMA2017SF-1951-TB-SPEC to CR), the NIH (R21AI148027 to CR) and the South African Medical Research Council (MRC-UFSP-1-IMPI-2 to MN). RJW is supported by the Francis Crick Institute, which receives funds from Cancer Research UK(FC00110218), Wellcome (FC00110218) and the UK Medical Research Council (FC00110218). RJW is also supported by Wellcome (203135), and NIH (U01/115940; U01/152103). HM is supported by National Research Foundation of South Africa, (Grant number: 129614), CIDRI-Africa Fellowship and in part by the Fogarty International Center of the National Institutes of Health (D43TW010559). DLB, KDMB, and AS are supported by the National Institute of Allergy and Infectious Diseases, National Institutes of Health, Division of Intramural Research.

## Competing interests

The authors declare that they have no competing interests associated with this publication.

## Notes

### Competing Interest Statement

The authors have declared no competing interest.

## REFERANCES

[1] Global HIV & AIDS statistics — Fact sheet, (n.d.). https://www.unaids.org/en/resources/fact-sheet (accessed January 31, 2022).

[2] S.K. Sharma, A. Mohan, Extrapulmonary tuberculosis., Indian J. Med. Res. 120 (2004) 316–353.

[3] A.A. Cagatay, Y. Caliskan, S. Aksoz, L. Gulec, S. Kucukoglu, Y. Cagatay, H. Berk, H. Ozsut, H. Eraksoy, S. Calangu, Extrapulmonary tuberculosis in immunocompetent adults, Scand. J. Infect. Dis. 36 (2004) 799–806. https://doi.org/10.1080/00365540410025339.

[4] R.W. Shafer, D.S. Kim, J.P. Weiss, J.M. Quale, Extrapulmonary tuberculosis in patients with human immunodeficiency virus infection, Medicine (Baltimore). 70 (1991) 384– 397. https://doi.org/10.1097/00005792-199111000-00004.

[5] H.L. Rieder, D.E. Snider, G.M. Cauthen, Extrapulmonary tuberculosis in the United States, Am. Rev. Respir. Dis. 141 (1990) 347–351. https://doi.org/10.1164/ajrccm/141.2.347.

[6] J.J. Noubiap, V.N. Agbor, A.L. Ndoadoumgue, J.R. Nkeck, A. Kamguia, U.F. Nyaga, M. Ntsekhe, Epidemiology of pericardial diseases in Africa: a systematic scoping review, Heart. 105 (2019) 180–188. https://doi.org/10.1136/heartjnl-2018-313922.

[7] P. Howlett, E. Du Bruyn, H. Morrison, I.C. Godsent, K.A. Wilkinson, M. Ntsekhe, R.J. Wilkinson, The immunopathogenesis of tuberculous pericarditis, Microbes Infect. 22 (2020) 172–181. https://doi.org/10.1016/j.micinf.2020.02.001.

[8] H. Reuter, L.J. Burgess, A.F. Doubell, Epidemiology of pericardial effusions at a large academic hospital in South Africa, Epidemiol. Infect. 133 (2005) 393–399. https://doi.org/10.1017/S0950268804003577.

[9] B.M. Mayosi, M. Ntsekhe, J. Bosch, S. Pandie, H. Jung, F. Gumedze, J. Pogue, L. Thabane, M. Smieja, V. Francis, L. Joldersma, K.M. Thomas, B. Thomas, A.A. Awotedu, N.P. Magula, D.P. Naidoo, A. Damasceno, A. Chitsa Banda, B. Brown, P. Manga, B. Kirenga, C. Mondo, P. Mntla, J.M. Tsitsi, F. Peters, M.R. Essop, J.B.W. Russell, J. Hakim, J. Matenga, A.F. Barasa, M.U. Sani, T. Olunuga, O. Ogah, V. Ansa, A. Aje, S. Danbauchi, D. Ojji, S. Yusuf, Prednisolone and Mycobacterium indicus pranii in Tuberculous Pericarditis, N. Engl. J. Med. 371 (2014) 1121–1130. https://doi.org/10.1056/NEJMoa1407380.

[10] B.M. Mayosi, C.S. Wiysonge, M. Ntsekhe, F. Gumedze, J.A. Volmink, G. Maartens, A. Aje, B.M. Thomas, K.M. Thomas, A.A. Awotedu, B. Thembela, P. Mntla, F. Maritz, K.N. Blackett, D.C. Nkouonlack, V.C. Burch, K. Rebe, A. Parrish, K. Sliwa, B.Z. Vezi, N. Alam, B.G. Brown, T. Gould, T. Visser, N.P. Magula, P.J. Commerford, Mortality in patients treated for tuberculous pericarditis in sub-Saharan Africa: original article, S. Afr. Med. J. 98 (2008) 36–40. https://doi.org/10.10520/EJC69118.

[11] C. Geldmacher, A. Zumla, M. Hoelscher, Interaction between HIV and Mycobacterium tuberculosis: HIV-1-induced CD4 T-cell depletion and the development of active tuberculosis, Curr. Opin. HIV Aids. 7 (2012) 268–275. https://doi.org/10.1097/coh.0b013e3283524e32.

[12] C. Riou, N. Strickland, A.P. Soares, B. Corleis, D.S. Kwon, E.J. Wherry, R.J. Wilkinson, W.A. Burgers, HIV Skews the Lineage-Defining Transcriptional Profile of Mycobacterium tuberculosis–Specific CD4+ T Cells, J. Immunol. 196 (2016) 3006–3018. https://doi.org/10.4049/jimmunol.1502094.

[13] J.M. Brenchley, M. Paiardini, K.S. Knox, A.I. Asher, B. Cervasi, T.E. Asher, P. Scheinberg, D.A. Price, C.A. Hage, L.M. Kholi, A. Khoruts, I. Frank, J. Else, T. Schacker, G. Silvestri, D.C. Douek, Differential Th17 CD4 T-cell depletion in pathogenic and nonpathogenic lentiviral infections, Blood. 112 (2008) 2826–2835. https://doi.org/10.1182/blood-2008-05-159301.

[14] S. Clark, E. Page, T. Ford, R. Metcalf, A. Pozniak, M. Nelson, D.C. Henderson, D. Asboe, F. Gotch, B.G. Gazzard, P. Kelleher, Reduced T(H)1/T(H)17 CD4 T-cell numbers are associated with impaired purified protein derivative-specific cytokine responses in patients with HIV-1 infection, J. Allergy Clin. Immunol. 128 (2011) 838-846.e5. https://doi.org/10.1016/j.jaci.2011.05.025.

[15] H. Mutavhatsindi, G.D. Van Der Spuy, S. Malherbe, J.S. Sutherland, A. Geluk, H. Mayanja Kizza, A.C. Crampin, D. Kassa, R. Howe, A. Mihret, J.A. Sheehama, E. Nepolo, G. Günther, H.M. Dockrell, P.L. Lam Corstjens, K. Stanley, G. Walzl, N.N. Chegou, Validation and optimisation of host immunological bio-signatures for a point-of-care test for TB disease, Front. Immunol. 12 (2021). https://doi.org/10.3389/fimmu.2021.607827.

[16] B.H. Chendi, H. Tveiten, C.I. Snyders, K. Tonby, S. Jenum, S.D. Nielsen, M. Hove-Skovsgaard, G. Walzl, N.N. Chegou, A.M. Dyrhol-Riise, CCL1 and IL-2Ra differentiate Tuberculosis disease from latent infection Irrespective of HIV infection in low TB burden countries, J. Infect. 83 (2021) 433–443. https://doi.org/10.1016/j.jinf.2021.07.036.

[17] N.N. Chegou, J.S. Sutherland, S. Malherbe, A.C. Crampin, P.L.A.M. Corstjens, A. Geluk, H. Mayanja-Kizza, A.G. Loxton, G. van der Spuy, K. Stanley, L.A. Kotzé, M. van der Vyver, I. Rosenkrands, M. Kidd, P.D. van Helden, H.M. Dockrell, T.H.M. Ottenhoff, S.H.E. Kaufmann, G. Walzl, Diagnostic performance of a seven-marker serum protein biosignature for the diagnosis of active TB disease in African primary healthcare clinic attendees with signs and symptoms suggestive of TB, Thorax. 71 (2016) 785–794. https://doi.org/10.1136/thoraxjnl-2015-207999.

[18] R. Jacobs, S. Malherbe, A.G. Loxton, K. Stanley, G. van der Spuy, G. Walzl, N.N. Chegou, Identification of novel host biomarkers in plasma as candidates for the immunodiagnosis of tuberculosis disease and monitoring of tuberculosis treatment response, Oncotarget. 7 (2016). https://doi.org/10.18632/oncotarget.11420.

[19] Q.-Y. Liu, F. Han, L.-P. Pan, H.-Y. Jia, Q. Li, Z.-D. Zhang, Inflammation responses in patients with pulmonary tuberculosis in an intensive care unit, Exp. Ther. Med. 15 (2018) 2719–2726. https://doi.org/10.3892/etm.2018.5775.

[20] O.A. Eribo, M.S. Leqheka, S.T. Malherbe, S. McAnda, K. Stanley, G.D. van der Spuy, G. Walzl, N.N. Chegou, Host urine immunological biomarkers as potential candidates for the diagnosis of tuberculosis, Int. J. Infect. Dis. 99 (2020) 473–481. https://doi.org/10.1016/j.ijid.2020.08.019.

[21] J.S. Sutherland, B.C. de Jong, D.J. Jeffries, I.M. Adetifa, M.O.C. Ota, Production of TNF-α, IL-12(p40) and IL-17 Can Discriminate between Active TB Disease and Latent Infection in a West African Cohort, PLOS ONE. 5 (2010) e12365. https://doi.org/10.1371/journal.pone.0012365.

[22] K. Matthews, A. Deffur, M. Ntsekhe, F. Syed, J.B.W. Russell, K. Tibazarwa, J. Wolske, J. Brink, B.M. Mayosi, R.J. Wilkinson, K.A. Wilkinson, A Compartmentalized Profibrotic Immune Response Characterizes Pericardial Tuberculosis, Irrespective of HIV-1 Infection, Am. J. Respir. Crit. Care Med. 192 (2015) 1518–1521. https://doi.org/10.1164/rccm.201504-0683LE.

[23] H. Reuter, L.J. Burgess, M.E. Carstens, A.F. Doubell, Characterization of the immunological features of tuberculous pericardial effusions in HIV positive and HIV negative patients in contrast with non-tuberculous effusions, Tuberculosis. 86 (2006) 125–133. https://doi.org/10.1016/j.tube.2005.08.018.

[24] Q. Yang, Y. Cai, W. Zhao, F. Wu, M. Zhang, K. Luo, Y. Zhang, H. Liu, B. Zhou, H. Kornfeld, X. Chen, IP-10 and MIG Are Compartmentalized at the Site of Disease during Pleural and Meningeal Tuberculosis and Are Decreased after Antituberculosis Treatment, Clin. Vaccine Immunol. 21 (2014) 1635–1644. https://doi.org/10.1128/CVI.00499-14.

[25] B.M. Mayosi, L.J. Burgess, A.F. Doubell, Tuberculous Pericarditis, Circulation. 112 (2005) 3608–3616. https://doi.org/10.1161/CIRCULATIONAHA.105.543066.

[26] W.A. Hanekom, J. Hughes, M. Mavinkurve, M. Mendillo, M. Watkins, H. Gamieldien, S.J. Gelderbloem, M. Sidibana, N. Mansoor, V. Davids, R.A. Murray, A. Hawkridge, P.A.J. Haslett, S. Ress, G.D. Hussey, G. Kaplan, Novel application of a whole blood intracellular cytokine detection assay to quantitate specific T-cell frequency in field studies, J. Immunol. Methods. 291 (2004) 185–195. https://doi.org/10.1016/j.jim.2004.06.010.

[27] C.S.L. Arlehamn, D.M. McKinney, C. Carpenter, S. Paul, V. Rozot, E. Makgotlho, Y. Gregg, M. van Rooyen, J.D. Ernst, M. Hatherill, W.A. Hanekom, B. Peters, T.J. Scriba, A. Sette, A Quantitative Analysis of Complexity of Human Pathogen-Specific CD4 T Cell Responses in Healthy M. tuberculosis Infected South Africans, PLOS Pathog. 12 (2016) e1005760. https://doi.org/10.1371/journal.ppat.1005760.

[28] R. Fluss, D. Faraggi, B. Reiser, Estimation of the Youden Index and its Associated Cutoff Point, Biom. J. 47 (2005) 458–472. https://doi.org/10.1002/bimj.200410135.

[29] J.M. Bland, D.G. Altman, Multiple significance tests: the Bonferroni method, BMJ. 310 (1995) 170. https://doi.org/10.1136/bmj.310.6973.170.

[30] D. Wang, X. Tong, L. Wang, S. Zhang, J. Huang, L. Zhang, H. Fan, The association between osteopontin and tuberculosis: A systematic review and meta-analysis, PLOS ONE. 15 (2020) e0242702. https://doi.org/10.1371/journal.pone.0242702.

[31] D.M. Muema, N.A. Akilimali, O.C. Ndumnego, S.S. Rasehlo, R. Durgiah, D.B.A. Ojwach, N. Ismail, M. Dong, A. Moodley, K.L. Dong, Z.M. Ndhlovu, J.M. Mabuka, B.D. Walker, J.K. Mann, T. Ndung’u, Association between the cytokine storm, immune cell dynamics, and viral replicative capacity in hyperacute HIV infection, BMC Med. 18 (2020) 81. https://doi.org/10.1186/s12916-020-01529-6.

[32] J.E. Teigler, L. Leyre, N. Chomont, B. Slike, N. Jian, M.A. Eller, N. Phanuphak, E. Kroon, S. Pinyakorn, L.A. Eller, M.L. Robb, J. Ananworanich, N.L. Michael, H. Streeck, S.J. Krebs, Distinct biomarker signatures in HIV acute infection associate with viral dynamics and reservoir size, JCI Insight. 3 (n.d.) e98420. https://doi.org/10.1172/jci.insight.98420.

[33] Á.H. Borges, J.L. O’Connor, A.N. Phillips, F.F. Rönsholt, S. Pett, M.J. Vjecha, M.A. French, J.D. Lundgren, Factors Associated With Plasma IL-6 Levels During HIV Infection, J. Infect. Dis. 212 (2015) 585–595. https://doi.org/10.1093/infdis/jiv123.

[34] L. Roberts, J.-A.S. Passmore, C. Williamson, F. Little, L.M. Bebell, K. Mlisana, W.A. Burgers, F. Van Loggerenberg, G. Walzl, J.F. DJOBA Siawaya, Q. ABDOOL Karim, S.S. ABDOOL Karim, Plasma cytokine levels during acute HIV-1 infection predict HIV disease progression, AIDS Lond. Engl. 24 (2010) 819–831. https://doi.org/10.1097/QAD.0b013e3283367836.

[35] R. Bunjun, A.P. Soares, N. Thawer, T.L. Müller, A. Kiravu, Z. Ginbot, B. Corleis, B.D. Murugan, D.S. Kwon, F. von Groote-Bidlingmaier, C. Riou, R.J. Wilkinson, G. Walzl, W.A. Burgers, Dysregulation of the Immune Environment in the Airways During HIV Infection, Front. Immunol. 12 (2021). https://www.frontiersin.org/article/10.3389/fimmu.2021.707355 (accessed April 12, 2022).

[36] G.P. Rizzardi, W. Barcellini, G. Tambussi, F. Lillo, M. Malnati, L. Perrin, A. Lazzarin, Plasma levels of soluble CD30, tumour necrosis factor (TNF)-alpha and TNF receptors during primary HIV-1 infection: correlation with HIV-1 RNA and the clinical outcome, AIDS Lond. Engl. 10 (1996) F45–50. https://doi.org/10.1097/00002030-199611000-00001.

[37] T. Adekambi, C.C. Ibegbu, S. Cagle, A.S. Kalokhe, Y.F. Wang, Y. Hu, C.L. Day, S.M. Ray, J. Rengarajan, Biomarkers on patient T cells diagnose active tuberculosis and monitor treatment response, J. Clin. Invest. 125 (2015) 1827–1838. https://doi.org/10.1172/JCI77990.

[38] C. Riou, N. Berkowitz, R. Goliath, W.A. Burgers, R.J. Wilkinson, Analysis of the Phenotype of Mycobacterium tuberculosis-Specific CD4+ T Cells to Discriminate Latent from Active Tuberculosis in HIV-Uninfected and HIV-Infected Individuals, Front. Immunol. 8 (2017). https://www.frontiersin.org/article/10.3389/fimmu.2017.00968 (accessed April 13, 2022).

[39] C. Riou, E. Du Bruyn, S. Ruzive, R.T. Goliath, C.S. Lindestam Arlehamn, A. Sette, A. Sher, D.L. Barber, R.J. Wilkinson, Disease extent and anti-tubercular treatment response correlates with Mycobacterium tuberculosis-specific CD4 T-cell phenotype regardless of HIV-1 status, Clin. Transl. Immunol. 9 (2020) e1176. https://doi.org/10.1002/cti2.1176.

[40] C.L. Vinhaes, D. Oliveira-de-Souza, P.S. Silveira-Mattos, B. Nogueira, R. Shi, W. Wei, X. Yuan, G. Zhang, Y. Cai, C.E. Barry, L.E. Via, K.F. Fukutani, B.B. Andrade, K.D. Mayer-Barber, Changes in Inflammatory Protein and Lipid Mediator Profiles Persist After Antitubercular Treatment of Pulmonary and Extrapulmonary Tuberculosis: A Prospective Cohort Study, Cytokine. 123 (2019) 154759. https://doi.org/10.1016/j.cyto.2019.154759.

[41] E. Du Bruyn, S. Ruzive, P. Howlett, A.J. Jacobs, C.S. Lindestam Arlehamn, A. Sette, A. Sher, K.D. Mayer-Barber, D.L. Barber, B. Mayosi, M. Ntsekhe, R.J. Wilkinson, C. Riou, Profile of Mycobacterium tuberculosis-specific CD4 T cells at the site of disease and blood in pericardial tuberculosis, BioRxiv. (2022) 2022.05.12.491749. https://doi.org/10.1101/2022.05.12.491749.

[42] S.G. Hoft, M.A. Sallin, K.D. Kauffman, S. Sakai, V.V. Ganusov, D.L. Barber, The Rate of CD4 T Cell Entry into the Lungs during Mycobacterium tuberculosis Infection Is Determined by Partial and Opposing Effects of Multiple Chemokine Receptors, Infect. Immun. 87 (2019) e00841–18. https://doi.org/10.1128/IAI.00841-18.

[43] P.S. Silveira-Mattos, B. Barreto-Duarte, B. Vasconcelos, K.F. Fukutani, C.L. Vinhaes, D. Oliveira-De-Souza, C.C. Ibegbu, M.C. Figueiredo, T.R. Sterling, J. Rengarajan, B.B. Andrade, Differential Expression of Activation Markers by Mycobacterium tuberculosis-specific CD4+ T Cell Distinguishes Extrapulmonary From Pulmonary Tuberculosis and Latent Infection, Clin. Infect. Dis. Off. Publ. Infect. Dis. Soc. Am. 71 (2020) 1905–1911. https://doi.org/10.1093/cid/ciz1070.

[44] World Health Organization, High priority target product profiles for new tuberculosis diagnostics: report of a consensus meeting, 28-29 April 2014, Geneva, Switzerland, World Health Organization, 2014. https://apps.who.int/iris/handle/10665/135617 (accessed June 3, 2022).

[45] D.A. Mitchison, Assessment of New Sterilizing Drugs for Treating Pulmonary Tuberculosis by Culture at 2 Months, Am. Rev. Respir. Dis. 147 (1993) 1062–1063. https://doi.org/10.1164/ajrccm/147.4.1062.

[46] R.S. Wallis, M. Pai, D. Menzies, T.M. Doherty, G. Walzl, M.D. Perkins, A. Zumla, Biomarkers and diagnostics for tuberculosis: progress, needs, and translation into practice, The Lancet. 375 (2010) 1920–1937. https://doi.org/10.1016/S0140-6736(10)60359-5.

[47] C. Riou, B.P. Peixoto, L. Roberts, K. Ronacher, G. Walzl, C. Manca, R. Rustomjee, T. Mthiyane, D. Fallows, C.M. Gray, G. Kaplan, Effect of Standard Tuberculosis Treatment on Plasma Cytokine Levels in Patients with Active Pulmonary Tuberculosis, PLOS ONE. 7 (2012) e36886. https://doi.org/10.1371/journal.pone.0036886.

[48] J.Y. Hong, H.J. Lee, S.Y. Kim, K.S. Chung, E.Y. Kim, J.Y. Jung, M.S. Park, Y.S. Kim, S.K. Kim, J. Chang, S.-N. Cho, Y.A. Kang, Efficacy of IP-10 as a biomarker for monitoring tuberculosis treatment, J. Infect. 68 (2014) 252–258. https://doi.org/10.1016/j.jinf.2013.09.033.

[49] C. Martins, A.C. de C. Gama, D. Valcarenghi, A.P. de B. Batschauer, C. Martins, A.C. de C. Gama, D. Valcarenghi, A.P. de B. Batschauer, Markers of acute-phase response in the treatment of pulmonary tuberculosis, J. Bras. Patol. E Med. Lab. 50 (2014) 428–433. https://doi.org/10.5935/1676-2444.20140052.

[50] B.H. Chendi, C.I. Snyders, K. Tonby, S. Jenum, M. Kidd, G. Walzl, N.N. Chegou, A.M. Dyrhol-Riise, A Plasma 5-Marker Host Biosignature Identifies Tuberculosis in High and Low Endemic Countries, Front. Immunol. 12 (2021). https://www.frontiersin.org/article/10.3389/fimmu.2021.608846 (accessed June 3, 2022).

[51] L. Liang, R. Shi, X. Liu, X. Yuan, S. Zheng, G. Zhang, W. Wang, J. Wang, K. England, L.E. Via, Y. Cai, L.C. Goldfeder, L.E. Dodd, C.E. Barry, R.Y. Chen, Interferon-gamma response to the treatment of active pulmonary and extra-pulmonary tuberculosis, Int. J. Tuberc. Lung Dis. 21 (2017) 1145–1149. https://doi.org/10.5588/ijtld.16.0880.

[52] N. Rockwood, D.L. Costa, E.P. Amaral, E. Du Bruyn, A. Kubler, L. Gil-Santana, K.F. Fukutani, C.A. Scanga, J.L. Flynn, S.H. Jackson, K.A. Wilkinson, W.R. Bishai, A. Sher, R.J. Wilkinson, B.B. Andrade, Mycobacterium tuberculosis induction of heme oxygenase-1 expression is dependent on oxidative stress and reflects treatment outcomes, Front. Immunol. 8 (2017). https://doi.org/10.3389/fimmu.2017.00542.

[53] R.S. Wallis, P. Kim, S. Cole, D. Hanna, B.B. Andrade, M. Maeurer, M. Schito, A. Zumla, Tuberculosis biomarkers discovery: developments, needs, and challenges, Lancet Infect. Dis. 13 (2013) 362–372. https://doi.org/10.1016/S1473-3099(13)70034-3.

[54] X. Bai, H. Li, Y. Yang, J. Zhang, Y. Liang, X. Wu, Cytokine and soluble adhesion molecule profiles and biomarkers for treatment monitoring in Re-treated smear-positive patients with pulmonary tuberculosis, Cytokine. 108 (2018) 9–16. https://doi.org/10.1016/j.cyto.2018.03.009.

[55] P. Miranda, L. Gil-Santana, M.G. Oliveira, E.D.D. Mesquita, E. Silva, A. Rauwerdink, F. Cobelens, M.M. Oliveira, B.B. Andrade, A. Kritski, Sustained elevated levels of C-reactive protein and ferritin in pulmonary tuberculosis patients remaining culture positive upon treatment initiation, PLOS ONE. 12 (2017) e0175278. https://doi.org/10.1371/journal.pone.0175278.

[56] G.B. Sigal, M.R. Segal, A. Mathew, L. Jarlsberg, M. Wang, S. Barbero, N. Small, K. Haynesworth, J.L. Davis, M. Weiner, W.C. Whitworth, J. Jacobs, J. Schorey, D.M. Lewinsohn, P. Nahid, Biomarkers of Tuberculosis Severity and Treatment Effect: A Directed Screen of 70 Host Markers in a Randomized Clinical Trial, EBioMedicine. 25 (2017) 112–121. https://doi.org/10.1016/j.ebiom.2017.10.018.

[57] A. Singanayagam, K. Manalan, D.W. Connell, J.D. Chalmers, S. Sridhar, A.I. Ritchie, A. Lalvani, M. Wickremasinghe, O.M. Kon, Evaluation of serum inflammatory biomarkers as predictors of treatment outcome in pulmonary tuberculosis, Int. J. Tuberc. Lung Dis. 20 (2016) 1653–1660. https://doi.org/10.5588/ijtld.16.0159.

[58] J. Nouhin, P. Pean, Y. Madec, M.F. Chevalier, C. Didier, L. Borand, F.-X. Blanc, D. Scott-Algara, D. Laureillard, L. Weiss, Interleukin-1 receptor antagonist, a biomarker of response to anti-TB treatment in HIV/TB co-infected patients, J. Infect. 74 (2017) 456– 465. https://doi.org/10.1016/j.jinf.2017.01.016.

[59] C. Schutz, D. Barr, B.B. Andrade, M. Shey, A. Ward, S. Janssen, R. Burton, K.A. Wilkinson, B. Sossen, K.F. Fukutani, M. Nicol, G. Maartens, R.J. Wilkinson, G. Meintjes, Clinical, microbiologic, and immunologic determinants of mortality in hospitalized patients with HIV-associated tuberculosis: A prospective cohort study, PLoS Med. 16 (2019) e1002840. https://doi.org/10.1371/journal.pmed.1002840.

