## Supplementery figures and tables for "Blood and site of disease inflammatory profiles differ in HIV-1-infected pericardial tuberculosis patients"

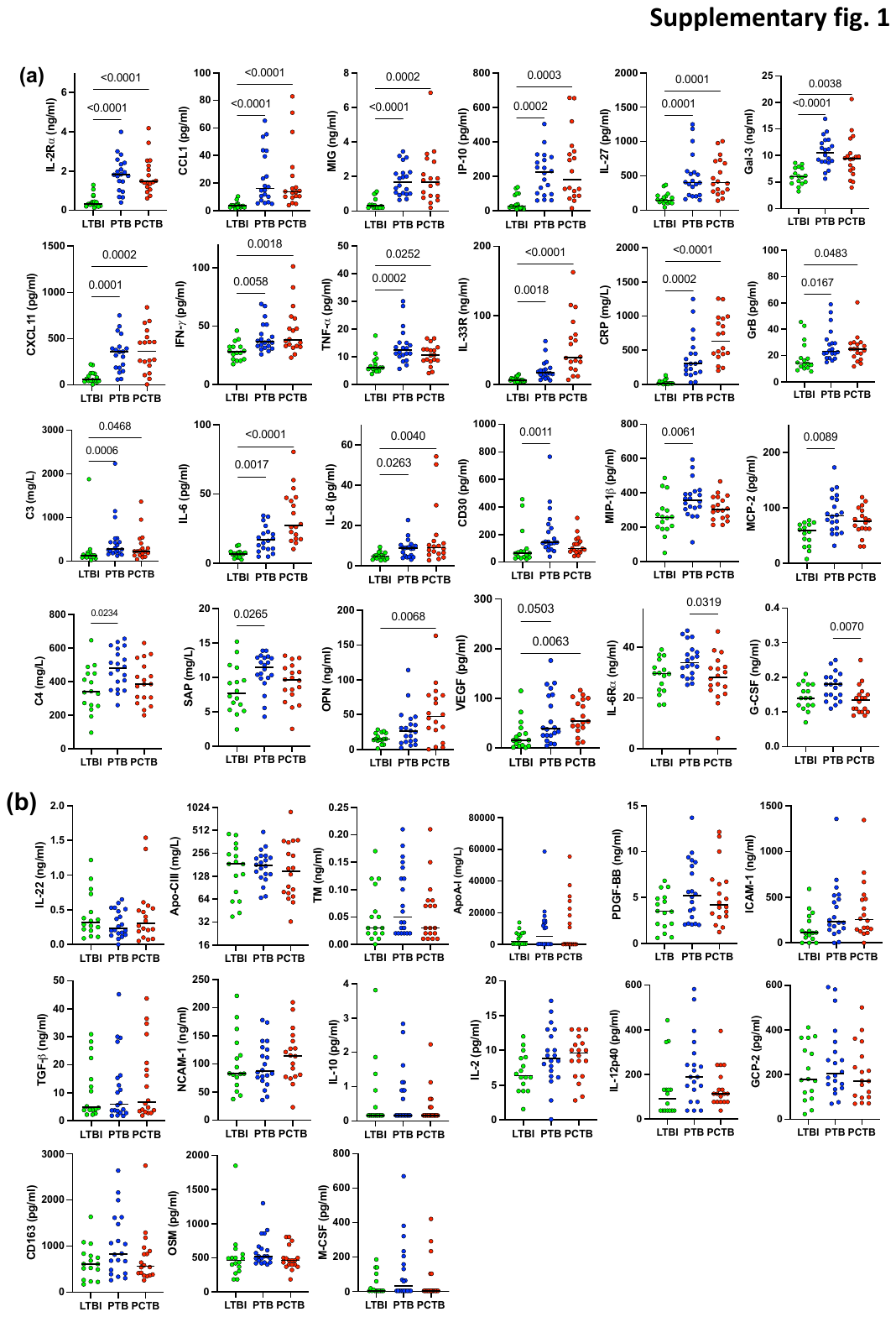

**Supplementary fig. 1.** **Scatter plots of the 39 analytes detected in plasma of participants with LTBI, PTB and PCTB.** **(a)** Analytes that are significantly elevated in diseased groups compared to LTBI. **(b)** Analytes that were not expressed differently between the three groups. Statistical comparisons were performed using a Kruskal-Wallis test, adjusted for multiple comparisons (Dunn’s test).

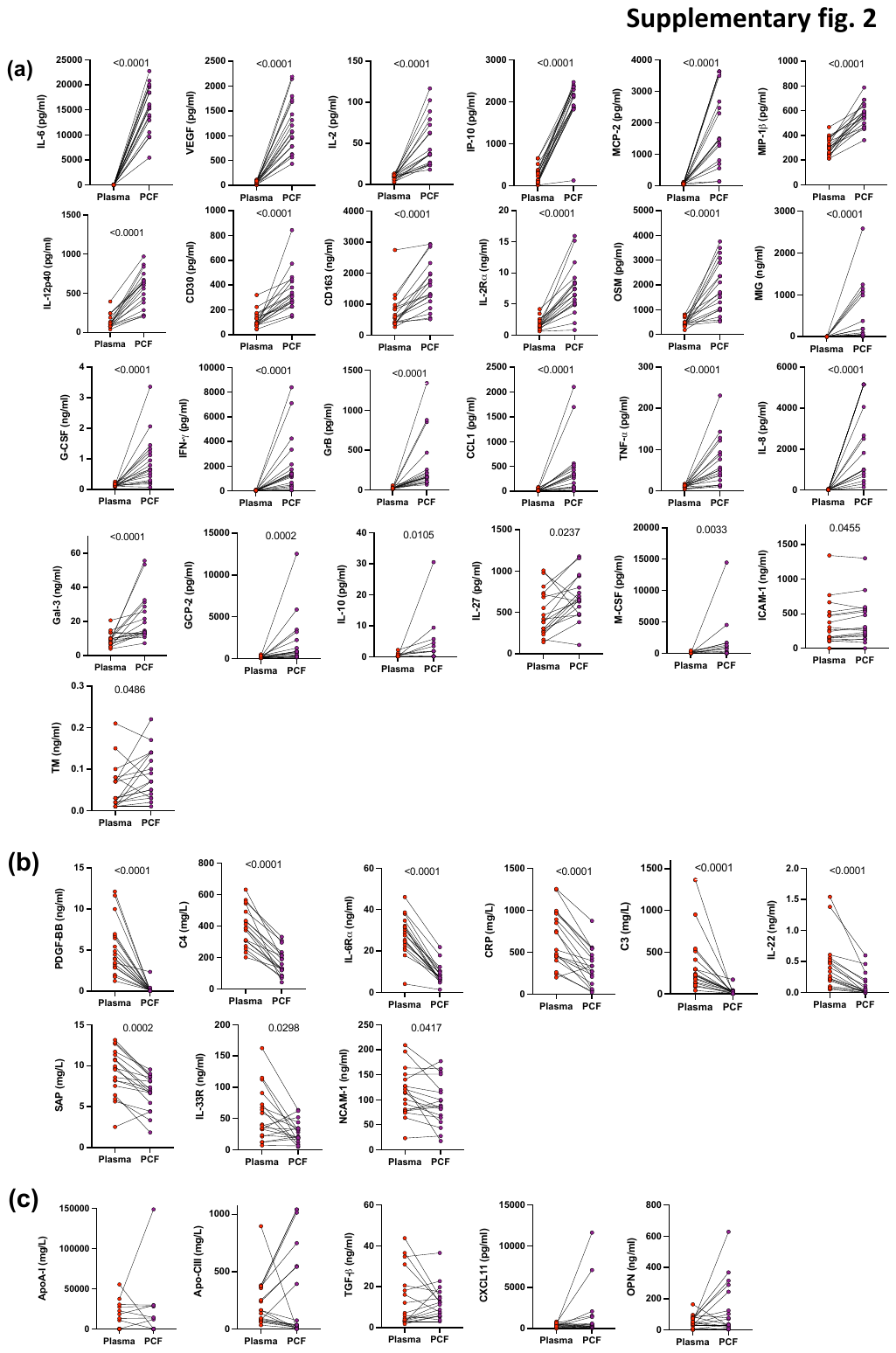

**Supplementary fig. 2.** **Baseline levels of analytes detected in Plasma and PCF of participants with PCTB.** **(a)** Analytes elevated in PCF compared to Plasma. **(b)** Analytes elevated in Plasma compared to PCF. **(c)** Analytes that showed no difference between Plasma and PCF. Statistical comparisons were performed using a Wilcoxon test and p-values were adjusted using the Bonferroni method.

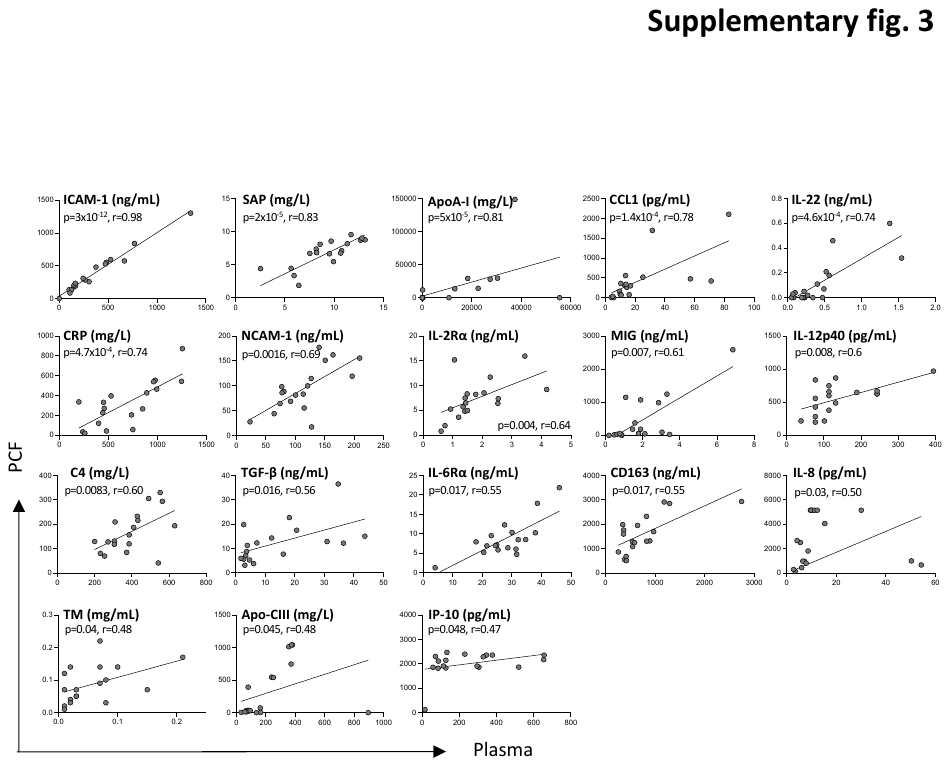

**Supplementary fig. 3.** **Univariate correlation of analytes detected in Plasma and PCF of participants with PCTB.** Analytes are arranged according to their correlation strength. Only analytes with significant associations are shown. The line indicates linear regression for statistically significant correlations. Correlations were tested by a two-tailed non-parametric Spearman rank test.

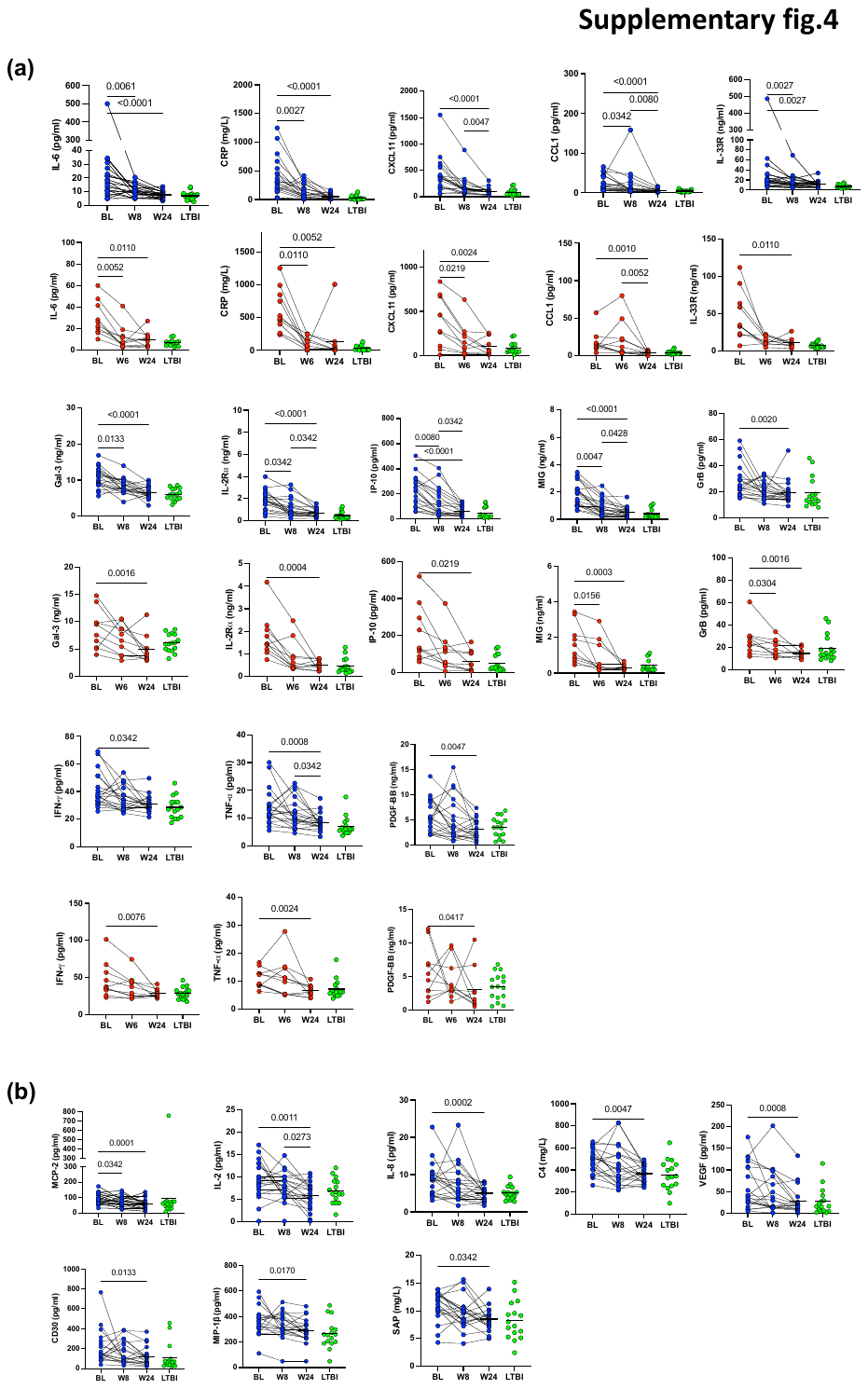

**Supplementary fig. 4.** **Longitudinal levels of analytes detected in plasma of participants with PTB and PCTB at Baseline, Week 6/8 post treatment initiation and at the end of treatment (Week 24). Week 24 was further compared and LTBI.** **(a)** Analytes that showed significant reduction with treatment in both PTB and PCTB. **(b)** Analytes that showed significant reduction with treatment in PTB only. Statistical comparisons were performed using a Friedman test, adjusted for multiple comparisons (Dunn’s test) for BL v W6/W8, BL v W24 and W6/W8 v W24 and the Mann-Whitney test to compare LTBI with W24, p-values were adjusted using the Bonferroni method.

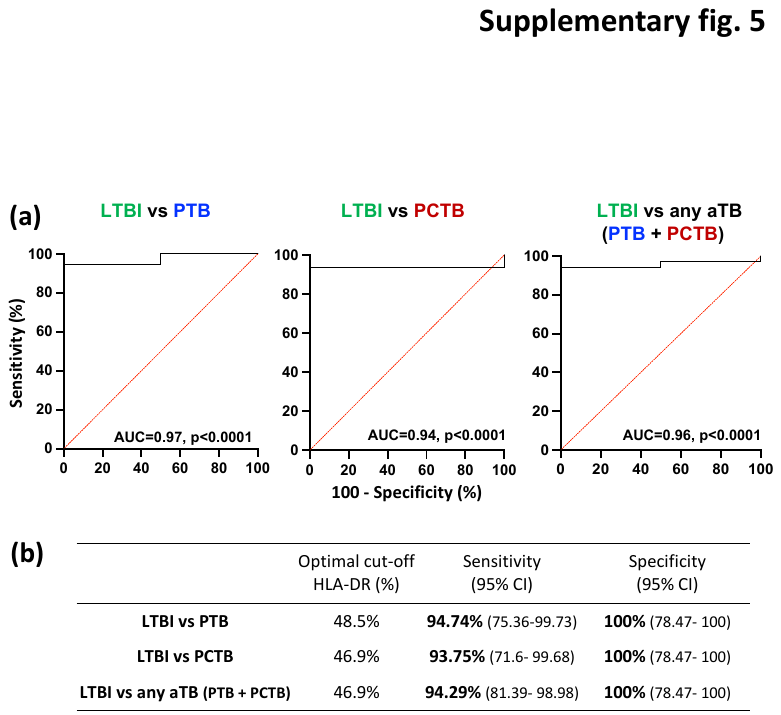

**Supplementary fig. 5.** **Ability of HLA-DR expression to discriminate LTBI from PTB, PCTB or any active TB (PTB + PCTB).** **(a)** Receiver operating characteristics (ROCs) curves for HLA-DR in discriminating LTBI from PTB, LTBI from PCTB and LTBI from any active TB (PTB + PCTB), respectively. **(b)** Corresponding sensitivity and specificity for each ROC curve at the optimal threshold of HLA-DR expression to distinguish between the groups.

**Supplementary table 1.** **Baseline median levels and interquartile ranges (IQR) of analytes detected in participants with PCTB, PTB and LTBI.**

|  |  |  |  |  | Adjusted P-Values | | | |
| --- | --- | --- | --- | --- | --- | --- | --- | --- |
| **Analytes** | **LTBI**  **n=16** | **PTB**  **n=20** | **PCTB**  **n=18** | **PTB & PCTB**  **n=38** | LTBI vs PTB | LTBI vs PCTB | PTB vs PCTB | LTBI vs All aTB |
| **Inflammation regulation** | | | | | | | | |
| GrB^†^ | 14.30  (11.07-25.48) | 23.22  (19-31.58) | 24.98  (18.71-29.57) | 24.37  (18.99-29.57) | **0.0167** | **0.0483** | >0.99 | **0.0057** |
| IL-2^†^ | 6.38  (4.41-8.983) | 8.87  (6.473-12.76) | 9.625  (6.27-11.25) | 9.25  (6.27-12) | 0.1331 | 0.1241 | >0.99 | **0.0360** |
| IL-8^†^ | 4.855  (3.693-6.235) | 8.73  (4.595-10.25) | 9.095  (5.918-13.09) | 8.73  (5.188-10.86) | **0.0263** | **0.0040** | >0.99 | **0.0017** |
| IL-12p40 ^†^ | 91.66  (38.48-131.7) | 188.8  (90.71-282.0) | 113.7  (77.04-202.3) | 140.4  (77.04-242.9) | 0.0594 | >0.99 | 0.4508 | 0.0942 |
| M-CSF^†^ | 4.13  (4.13-92.75) | 32  (4.13-19 3.8) | 4.13  (4.13-103.7) | 4.13  (4.13-137.8) | >0.99 | >0.99 | 0.5674 | 0.8245 |
| TNF-⍺^†^ | 6.09  (5.158-8.313) | 12.38  (10.25-15.12) | 10.57  (8.570-12.88) | 11.44  (8.680-14.37) | **0.0002** | **0.0252** | 0.491 | **<0.0001** |
| TGF-β^§^ | 4.865  (3.788-20.51) | 5.9  (2.965-15.94) | 6.645  (3.330-23.21) | 6.195  (3.330-18.79) | >0.99 | >0.99 | >0.99 | 0.9510 |
| C3^‡^ | 121.9  (85.9-188.5) | 284.8  (188.6-499.9) | 225.4  (137.1-435.4) | 248.5  (180.8-454.6) | **0.0006** | **0.0468** | 0.5989 | **0.0009** |
| C4^‡^ | 339.8  (256.9-448.7) | 479.8  (365.4-588.4) | 385.2  (297.5-503.4) | 430.7  (342.2-544.6) | **0.0243** | >0.99 | 0.1963 | 0.0837 |
| CRP^‡^ | 12.11  (10.88-41.73) | 301  (167.4-518.2) | 631.1  (432.2-965.1) | 447.9  (235.0-810.6) | **0.0002** | **<0.0001** | 0.1408 | **<0.0001** |
| SAP^‡^ | 7.64  (5.48-10.95) | 11.48  (9.86-12.85) | 9.61  (7.25-11.43) | 10.46  (8.21-12.69) | **0.0253** | >0.99 | 0.2256 | 0.0695 |
| IL-22^§^ | 0.32  (0.21-0.67) | 0.24  (0.15-0.50) | 0.31  (0.17-0.53) | 0.27  (0.16-0.51) | >0.99 | >0.99 | >0.99 | 0.5832 |
| Gal-3^§^ | 5.99  (4.80-7.65) | 10.56  (8.84-12.76) | 9.44  (7.01-11.11) | 9.62  (8.01-12.45) | **<0.0001** | **0.0038** | 0.7957 | **<0.0001** |
| ICAM-1^§^ | 111.6  (47.92-281.0) | 235.5  (147.3-508.6) | 254.4  (141.7-492.3) | 246.5  (146.1-490.4) | 0.0548 | 0.0647 | >0.99 | **0.0144** |
| NCAM-1^§^ | 82.79  (66.01-132.0) | 87.06  (71.61-128.5) | 114.3  (77.44-143.1) | 99.97  (76.82-132.2) | >0.99 | >0.99 | 0.9726 | 0.7161 |
| G-CSF^§^ | 0.140  (0.12-0.18) | 0.18  (0.14-0.21) | 0.14  (0.1-0.15) | 0.15  (0.12-0.19) | 0.1429 | >0.99 | **0.0104** | 0.5832 |
| IFN-γ ^†^ | 27.81  (21.51-32.20) | 36.61  (31.82-49.34) | 38.03  (32.73-57.08) | 37.35  (32.73-51.38) | **0.0058** | **0.0018** | >0.99 | **0.0003** |
| IL-6^†^ | 6.885  (4.62-8.163) | 17.13  (10.49-26.05) | 27.36  (17.72-47.42) | 21.28  (14.35-34.89) | **0.0017** | **<0.0001** | 0.0980 | **<0.0001** |
| IL-10^†^ | 0.16  (0.16-0.7675) | 0.16  (0.16-1.07) | 0.16  (0.16-0.46) | 0.16  (0.16-0.89) | >0.99 | >0.99 | 0.8161 | 0.8234 |
| IL-27^†^ | 149.7  (109.4-211.5) | 406.1  (223.2-565.8) | 405.6  (270.7-719.6) | 406.1  (253.0-691.7) | **0.0001** | **0.0001** | >0.99 | **<0.0001** |
| VEGF^†^ | 15.86  (4.64-38) | 39.4  (25.48-106.9) | 55.05  (39.92-98.64) | 48.24  (25.71-101.2) | 0.0503 | **0.0063** | >0.99 | **0.0039** |
| **Chemokines** | | | | | | | | |
| MIG^§^ | 0.31  (0.19-0.6) | 1.69  (1.02-2.24) | 1.69  (0.85-2.92) | 1.69  (0.96-2.4) | **<0.0001** | **0.0002** | >0.99 | **<0.0001** |
| MCP-2^†^ | 58.77  (33.53-72.68) | 85.79  (61.34-117) | 75.46  (62.2-96.47) | 79.1  (62.2-102.6) | **0.0089** | 0.1739 | 0.8660 | **0.0098** |
| GCP-2^†^ | 177.8  (99.72-351.2) | 203.8  (144-335.6) | 171.8  (98.85-298.1) | 191.3  (121.1-312.9) | >0.99 | >0.99 | 0.8647 | 0.8866 |
| CXCL11^†^ | 58.34  (43.09-125.5) | 359.2  (186.4-519.7) | 365.9  (189.2-535.3) | 359.2  (187.5-526.6) | **0.0001** | **0.0002** | >0.99 | **<0.0001** |
| MIP-1β^†^ | 256.8  (191.9-305.6 | 357.9  (306.3-404.8) | 302.8  (251.2-375.8) | 324.4  (272.6-388.5) | **0.0061** | 0.4046 | 0.3271 | **0.0144** |
| CCL1^†^ | 3.50  (2.3-4.29) | 16.01  (8.01-42.45) | 13.69  (9.89-26.77) | 14.45  (9.89-33.48) | **<0.0001** | **<0.0001** | >0.99 | **<0.0001** |
| IP-10^†^ | 27.53  (19.74-98.70) | 224.9  (98.06-308.1) | 181.7  (87.29-353.9) | 224.5  (94.93-325.1) | **0.0002** | **0.0003** | >0.99 | **<0.0001** |
| **Fibrosis regulation** | | | | | | | | |
| OSM^†^ | 460.8  (319.5-537.5) | 521.1  (437.5-656) | 460.9  (391.5-546.5) | 491.2  (417.5-640.7) | 0.1445 | >0.99 | 0.1877 | 0.2939 |
| IL-33R^§^ | 6.48  (4.43-8.9) | 17.25  (11.78-28.89) | 39.25  (22.52-76.88) | 23.56  (14.48-59.6) | **0.0018** | **<0.0001** | 0.0957 | **0.0003** |
| OPN^§^ | 14.83  (12.46-23.52) | 25.9  (11.05-41.96) | 47.4  (23.11-77.78) | 31.49  (11.6-55.53) | 0.3743 | **0.0068** | 0.3009 | **0.0173** |
| PDGF-BB^§^ | 3.49  (1.93-4.97) | 5.23  (2.34-8.74) | 4.22  (2.92-6.78) | 4.69  (2.86-7.82) | 0.1829 | 0.5483 | >0.99 | 0.0999 |
| TM^§^ | 0.03  (0.02-0.098) | 0.05  (0.02-0.14) | 0.03  (0.02-0.08) | 0.04  (0.02-0.11) | 0.5632 | >0.99 | 0.6208 | 0.5364 |
| **Chemokine and protein receptors** | | | | | | | | |
| CD163^§^ | 614.5  (323.5-827.2) | 829.4  (428.5-1582) | 568.2  (382.2-915.4) | 691.9  (405.2-1132) | 0.3216 | >0.99 | 0.5708 | 0.3533 |
| IL-6R⍺^§^ | 29.51  (22.89-34.3) | 33.86  (28.81-38.72) | 28.10  (22.66-32.79) | 31.74  (25.53-37.25) | 0.0589 | >0.99 | **0.0319** | 0.2840 |
| CD30^†^ | 64.01  (35.13-92.94) | 145.3  (110.4-268.5) | 97.57  (75.11-154.3) | 133.7  (85.46-186.2) | **0.0011** | 0.2352 | 0.2100 | **0.0041** |
| IL-2R⍺^§^ | 0.33  (0.24-0.66) | 1.82  (1.13-2.4) | 1.47  (1.165-2.34) | 1.75  (1.165-2.36) | **<0.0001** | **<0.0001** | >0.99 | **<0.0001** |
| **Apolipoproteins** | | | | | | | | |
| ApoA-I^‡^ | 2050  (364.2-7118) | 5264  (175-14620) | 228.2  (104.1-23813) | 263.4  (153.2-15722) | >0.99 | >0.99 | >0.99 | 0.9510 |
| Apo-CIII^‡^ | 185.5  (64.7-284.3) | 177.9  (126.7-234.4) | 148.7  (75.01-359.8) | 163.6  (89.71-241.8) | >0.99 | >0.99 | >0.99 | 0.9510 |

Statistical comparisons were performed using a Kruskal-Wallis test adjusted for multiple comparisons (Dunn’s test) for LTBI vs PTB, LTBI vs PCTB and PTB vs PCTB and the Mann-Whitney test to compare LTBI and aTB (PCTB & PTB) with p-values adjusted using the Bonferroni method.

^†^: Values are shown in pg/ml, ^‡^: Values are shown in mg/L ^§^: Values are shown in ng/ml.

Abbreviations. LTBI: Latent tuberculosis infection, PTB: Pulmonary Tuberculosis, PCTB, Pericardial tuberculosis, aTB: Active tuberculosis, Apo: Apolipoprotein, C3: Complement component 3, C4: Complement component 4, CRP: C reactive protein, SAP: Serum amyloid protein, IL: Interleukin, PDGF: Platelet-derived growth factor, MCP: Monocyte chemotactic protein, ICAM: Intercellular adhesion molecule, NCAM: Neural cell adhesion molecule, IP-10: Interferon γ-induced protein 10 kDa, OPN: Osteopontin, CD: Cluster of differentiation, MIG: Monokine induced by gamma interferon, G-CSF: Granulocyte colony-stimulating factor, IFN: Interferon, OSM: Oncostatin M, VEGF: Vascular endothelial growth factor, MIP: Macrophage inflammatory protein, GCP-2: granulocyte chemotactic protein 2, CXCL11: C-X-C motif chemokine 11, CCL1: C-C Motif Chemokine Ligand 1, M-CSF: Macrophage colony-stimulating factor, TM: Thrombomodulin, TNF-⍺: Tumour necrosis factor alpha and TGF-β: Transforming growth factor beta.

**Supplementary table 2. Baseline median levels and interquartile ranges (IQR) of analytes detected in Plasma and PCF of participants with PCTB.**

| **Analytes** | **Plasma (n=18)** | **PCF (n=18)** | **Adjusted P-values** |
| --- | --- | --- | --- |
| **Inflammation regulation** | | | |
| GrB (pg/mL) | 24.98 (18.71-29.57) | 168.7 (120.8-311.5) | **<0.0001** |
| IL-2 (pg/mL) | 9.63 (6.27-11.25) | 37.05 (25.23-74.25) | **<0.0001** |
| IL-8 (pg/mL) | 9.10 (5.92-13.09) | 2163 (778.1-5155) | **<0.0001** |
| IL-12p40 (pg/mL) | 113.7 (77.04-202.3) | 613.0 (356.9-691.1) | **<0.0001** |
| M-CSF (pg/mL) | 4.13 (4.13-103.7) | 214.8 (4.130-1351) | **0.0033** |
| TNF-⍺ (pg/mL) | 10.57 (8.57-12.88) | 50.76 (22.9-100.6) | **<0.0001** |
| TGF-β (ng/mL) | 6.65 (3.33-23.21) | 11.7 (5.89-15.69) | 0.7337 |
| C3 (mg/L) | 225.4 (137.1-435.4) | 17.29 (6.143-30.77) | **<0.0001** |
| C4 (mg/L) | 385.2 (297.5-503.4) | 144.5 (109.9-220.4) | **<0.0001** |
| CRP (mg/L) | 631.1 (432.2-965.1) | 301.8 (106.5-484.6) | **<0.0001** |
| SAP (mg/L) | 9.60 (7.25-11.43) | 6.97 (5.21-8.59) | **0.0002** |
| IL-22 (ng/mL) | 0.31 (0.17-0.53) | 0.04 (0-0.19) | **<0.0001** |
| Gal-3 (ng/mL) | 9.44 (7.01-11.11) | 14.37 (12.71-29.31) | **<0.0001** |
| ICAM-1 (ng/mL) | 254.4 (141.7-492.3) | 269.7 (165.4-558.4) | **0.0455** |
| NCAM-1 (ng/mL) | 114.3 (77.44-143.1) | 87.11 (62.06-127.2) | **0.0417** |
| G-CSF (ng/mL) | 0.14 (0.1-0.15) | 0.66 (0.29-1.26) | **<0.0001** |
| IFN- γ (pg/mL) | 38.03 (32.73-57.08) | 1336 (455.8-2452) | **<0.0001** |
| IL-6 (pg/mL) | 27.36 (17.72-47.42) | 15571 (12341-19607) | **<0.0001** |
| IL-10 (pg/mL) | 0.16 (0.16-0.46) | 0.16 (0.16-3.715) | **0.0105** |
| IL-27 (pg/mL) | 405.6 (270.7-719.6) | 652.6 (477.8-835.3) | **0.0237** |
| VEGF (pg/mL) | 55.05 (39.92-98.64) | 1080 (747.6-1695) | **<0.0001** |
| **Chemokines** | | | |
| MIG (ng/mL) | 1.69 (0.85-2.9) | 76.20 (25.66-1009) | **<0.0001** |
| MCP-2 (pg/mL) | 75.46 (62.2-96.47) | 1493 (782.5-3514) | **<0.0001** |
| GCP-2 (pg/mL) | 171.8 (98.85-298.1) | 789.8 (283-2470) | **0.0002** |
| CXCL11 (pg/mL) | 365.9 (189.2-535.3) | 329.6 (177.4-1454) | 0.2479 |
| MIP-1β (pg/mL) | 302.8 (251.2-375.8) | 573.5 (492.4-638.9) | **<0.0001** |
| CCL1 (pg/mL) | 13.69 (9.89-26.77) | 313.8 (77.73-490.9) | **<0.0001** |
| IP-10 (pg/mL) | 181.7 (87.29-353.9) | 2131 (1867-2360) | **<0.0001** |
| **Fibrosis** **regulation** | | | |
| OSM (pg/mL) | 460.9 (391.5-546.5) | 1674 (1010-2943) | **<0.0001** |
| IL-33R (ng/mL) | 39.25 (22.52-76.88) | 24.52 (16.04-36.78) | **0.0298** |
| OPN (ng/mL) | 47.40 (23.11-77.78) | 53.75 (20.01-255.4) | 0.0907 |
| PDGF-BB (ng/mL) | 4.215 (2.920-6.778) | 0.08 (0.0475-0.1125) | **<0.0001** |
| TM (ng/mL) | 0.03 (0.0175-0.08) | 0.07 (0.0375-0.14) | **0.0486** |
| **Chemokine and protein receptors** | | | |
| CD163 (ng/mL) | 568.2 (382.2-915.4) | 1462 (1033-2072) | **<0.0001** |
| IL-6R⍺ (ng/mL) | 28.1 (22.66-32.79) | 7.59 (6.058-10.32) | **<0.0001** |
| CD30 (pg/mL) | 97.57 (75.11-154.3) | 309.4 (250.2-434.4) | **<0.0001** |
| IL-2R⍺ (ng/mL) | 1.47 (1.17-2.34) | 6.94 (4.94-8.70) | **<0.0001** |
| **Apolipoproteins** | | | |
| ApoA-I (mg/L) | 228.2 (104.1-23813) | 137.3 (78.28-17790) | 0.7204 |
| Apo-CIII (mg/L) | 148.7 (75.01-359.8) | 34.87 (11.8-597.5) | 0.6424 |

Statistical comparisons were performed using a Wilcoxon test and p-values were adjusted using the Bonferroni method.

Abbreviations. Apo: Apolipoprotein, C3: Complement component 3, C4: Complement component 4, CRP: C reactive protein, SAP: Serum amyloid protein, IL: Interleukin, TM: Thrombomodulin, PDGF: Platelet-derived growth factor, MCP: monocyte chemotactic protein, ICAM: Intercellular adhesion molecule, NCAM: Neural cell adhesion molecule, IP-10: Interferon γ-induced protein 10 kDa, OPN: Osteopontin, CD: Cluster of differentiation, MIG: Monokine induced by interferon gamma, G-CSF: Granulocyte colony-stimulating factor, IFN-γ: Interferon-gamma, OSM: Oncostatin M, VEGF: Vascular endothelial growth factor, MIP: Macrophage inflammatory protein, GCP-2: Granulocyte chemotactic protein 2, CXCL: C-X-C motif ligand, CCL: C-C motif ligand, M-CSF: Macrophage colony-stimulating factor, TNF-⍺: Tumour necrosis factor alpha, and TGF-β: Transforming growth factor beta.

**Supplementary table 3. Longitudinal median levels and interquartile ranges (IQR) of analytes detected in participants with PCTB and PTB at Baseline, Week 6/8 post treatment initiation and at the end of treatment (Week 24).**

|  | LTBI | PTB | | | P-Values | | | | PCTB | | | P-Values | | | |
| --- | --- | --- | --- | --- | --- | --- | --- | --- | --- | --- | --- | --- | --- | --- | --- |
| Analytes | **Baseline**  **n=16** | **Baseline**  **n=20** | **Week 8**  **n=20** | **Week 24**  **n=20** | BL vs W8 | BL vs W24 | W8 vs W24 | LTBI vs W24 | **Baseline**  **n=10** | **Week 6**  **n=10** | **Week 24**  **n=10** | BL vs W6 | BL vs W24 | W6 vs W24 | LTBI vs W24 |
| GrB^†^ | 14.30  (11.07-25.48) | 23.22  (19.0-31.58) | 19.26  (15.99-25.55) | 17.37  (14.71-20.09) | 0.207 | **0.002** | 0.342 | 0.5489 | 23.51  (16.27-29.57) | 16.19  (12.59-25.37) | 13.78  (10.69-17.3) | **0.030** | **0.0016** | >0.99 | 0.975 |
| IL-2^†^ | 6.38  (4.41-8.98) | 8.87  (6.47-12.76) | 7.8  (5.55-9.81) | 5.73  (3.59-7.74) | >0.99 | **0.001** | **0.027** | 0.6425 | 10.02  (7.94-12.25) | 7.04  (4.84-12.25) | 7.16  (5.54-9.25) | 0.1325 | 0.1325 | >0.99 | 0.990 |
| IL-8^†^ | 4.86  (3.70-6.24) | 8.73  (4.59-10.25) | 5.91  (3.49-9.99) | 5.21  (3.39-6.13) | 0.291 | **0.0002** | 0.0531 | 0.9146 | 7.89  (4.01-16.76) | 5.65  (3.78-15.51) | 6.27  (3.33-9.43) | >0.99 | 0.0566 | 0.2806 | 0.975 |
| IL-12p40 ^†^ | 91.66  (38.48-131.7) | 188.8  (90.71-282.0) | 131.7  (38.48-249.1) | 113.7  (46.28-174.5) | >0.99 | 0.1733 | 0.4642 | 0.7276 | 95.35  (77.04-202.3) | 100.7  (38.48-268.5) | 69.66  (61.87-136.1) | >0.99 | 0.6563 | 0.7907 | 0.990 |
| M-CSF^†^ | 4.13  (4.13-92.75) | 32.0  (4.13-193.8) | 4.13  (4.13-66.32) | 4.13  (4.13-74.99) | 0.707 | >0.99 | >0.99 | 0.9481 | 4.13  (4.13-103.7) | 45.04  (4.13-147.0) | 15.42  (4.13-92.63) | >0.99 | >0.99 | >0.99 | 0.990 |
| TNF-α^†^ | 6.09  (5.16-8.31) | 12.38  (10.25-15.12) | 9.96  (7.76-14.43) | 7.81  (7.12-9.53) | 0.8051 | **0.0008** | **0.0342** | 0.1577 | 10.68  (8.57-13.44) | 10.46  (5.33-14.79) | 6.61  (4.76-8.19) | 0.5391 | **0.0024** | 0.1325 | 0.990 |
| TGF-β^§^ | 4.87  (3.79-20.51) | 5.9  (2.96-15.94) | 5.18  (3.30-12.05) | 4.47  (2.71-8.95) | >0.99 | >0.99 | >0.99 | 0.5489 | 16.13  (3.73-35.59) | 11.55  (3.57-18.34) | 4.24  (3.80-9.04) | >0.99 | >0.99 | >0.99 | 0.990 |
| C3^‡^ | 121.9  (85.9-188.5) | 284.8  (188.6-499.9) | 306.6  (154.3-496.9) | 245.1  (154.3-306.1) | >0.99 | 0.6177 | 0.3415 | 0.0874 | 202.2  (119.7-323.3) | 242.7  (139.1-666.3) | 113.8  (67.98-228.8) | >0.99 | 0.2209 | 0.0760 | 0.990 |
| C4^‡^ | 339.8  (256.9-448.7) | 479.8  (365.4-588.4) | 395.7  (284.1-540.9) | 368.0  (296.7-443.8) | 0.1195 | **0.0047** | 0.8051 | 0.7669 | 407.3  (261.7-544.6) | 349.9  (268.7-429.5) | 277.3  (252.2-456.5) | >0.99 | >0.99 | >0.99 | 0.990 |
| CRP^‡^ | 12.11  (10.88-41.73) | 301.0  (167.4-518.2) | 87.15  (29.93-252.5) | 21.47  (12.18-74.6) | **0.0027** | **<0.0001** | 0.1733 | 0.4473 | 503.7  (365.1-884.6) | 63.70  (13.64-188.4) | 20.77  (11.14-87.52) | **0.011** | **0.0052** | >0.99 | 0.975 |
| SAP^‡^ | 7.64  (5.48-10.95) | 11.48  (9.86-12.85) | 9.57  (7.92-10.46) | 8.53  (7.3-9.39) | 0.2460 | **0.0342** | >0.99 | 0.7070 | 9.08  (5.85-12.71) | 9.73  (5.62-11.58) | 8.08  (6.06-8.89) | >0.99 | 0.7907 | >0.99 | 0.990 |
| IL-22^§^ | 0.32  (0.21-0.67) | 0.24  (0.15-0.50) | 0.32  (0.13-0.42) | 0.25  (0.143-0.59) | >0.99 | 0.7070 | >0.99 | 0.7070 | 0.25  (0.095-0.82) | 0.47  (0.35-1.2) | 0.32  (0.22-0.55) | **0.0156** | >0.99 | **0.0304** | 0.990 |
| Gal-3^§^ | 5.99  (4.80-7.65) | 10.56  (8.84-12.76) | 8.04  (7.02-9.68) | 6.23  (5.52-8.06) | **0.0133** | **<0.0001** | 0.0531 | 0.6271 | 8.52  (5.18-10.77) | 7.11  (3.79-9.010) | 4.0  (3.22-5.69) | 0.1720 | **0.0016** | 0.3526 | 0.753 |
| ICAM-1^§^ | 111.6  (47.92-281.0) | 235.5  (147.3-508.6) | 257.7  (157.2-526.6) | 264.7  (157.5-427.0) | >0.99 | >0.99 | 0.3415 | 0.0969 | 274.6  (150.3-527.5) | 207.3  (158.1-876.9) | 260.0  (143.7-700.0) | >0.99 | >0.99 | >0.99 | 0.753 |
| NCAM-1^§^ | 82.79  (66.01-132.0) | 87.06  (71.61-128.5) | 101.9  (79.87-165.8) | 117.4  (72.51-178.9) | 0.4642 | 0.0531 | >0.99 | 0.5489 | 107.0  (76.70-154.6) | 122.4  (90.52-168.4) | 141.7  (105.5-182.7) | 0.2209 | **0.0110** | 0.7907 | 0.753 |
| G-CSF^§^ | 0.14  (0.12-0.18) | 0.18  (0.14-0.21) | 0.19  (0.14-0.2) | 0.17  (0.14-0.23) | >0.99 | >0.99 | >0.99 | 0.4473 | 0.12  (0.10-0.15) | 0.17  (0.16-0.20) | 0.16  (0.13-0.22) | 0.2209 | 0.3526 | >0.99 | 0.975 |
| IFN-γ^†^ | 27.81  (21.51-32.2) | 36.61  (31.82-49.34) | 33.07  (27.51-41.32) | 29.39  (27.51-33.50) | 0.6177 | 0.0342 | 0.6177 | 0.5489 | 36.56  (31.12-59.33) | 33.08  (23.62-44.81) | 26.38  (24.89-31.99) | 0.0566 | **0.0076** | >0.99 | 0.990 |
| IL-6^†^ | 6.89  (4.62-8.16) | 17.13  (10.49-26.05) | 10.27  (7.27-13.26) | 7.95  (6.22-9.55) | **0.0061** | **<0.0001** | 0.4642 | 0.5489 | 24.54  (17.40-43.17) | 7.09  (4.14-14.68) | 9.89  (3.53-12.66) | **0.0052** | **0.0110** | >0.99 | 0.975 |
| IL-10^†^ | 0.16  (0.16-0.77) | 0.16  (0.16-1.07) | 0.40  (0.16-1.37) | 0.16  (0.16-0.64) | >0.99 | >0.99 | >0.99 | 0.7070 | 0.16  (0.16-0.46) | 0.64  (0.16-1.01) | 0.64  (0.16-1.13) | 0.2806 | 0.1325 | >0.99 | 0.975 |
| IL-27^†^ | 149.7  (109.4-211.5) | 406.1  (223.2-565.8) | 400.6  (210.2-589.5) | 371.9  (170.9-475.6) | 0.1733 | 0.1733 | >0.99 | **0.0273** | 405.6  (286.8-586.1) | 338.3  (198.6-475.3) | 214.5  (133.1-350.4) | >0.99 | 0.1325 | 0.5391 | 0.894 |
| VEGF^†^ | 15.86  (4.64-38.00) | 39.40  (25.48-106.9) | 22.68  (12.28-80.40) | 17.72  (9.05-24.71) | 0.1195 | **0.0008** | 0.3415 | 0.9481 | 59.11  (44.36-104.4) | 31.73  (13.69-76.9) | 19.30  (14.50-35.87) | 0.5391 | 0.1325 | >0.99 | 0.975 |
| MIG^§^ | 0.305  (0.19-0.60) | 1.685  (1.018-2.238) | 0.695  (0.37-1.303) | 0.43  (0.19-0.72) | **0.0047** | **<0.0001** | **0.0428** | 0.7070 | 1.34  (0.732-2.42) | 0.345  (0.19-1.663) | 0.19  (0.19-0.395) | **0.0156** | **0.0003** | 0.7907 | 0.975 |
| MCP-2^†^ | 58.77  (33.53-72.68) | 85.79  (61.34-117.0) | 67.32  (52.85-94.16) | 56.87  (35.20-71.04) | **0.0342** | **0.0001** | 0.3415 | 0.9686 | 78.67  (61.37-87.98) | 72.34  (41.11-79.14) | 42.22  (25.38-71.86) | >0.99 | 0.7907 | >0.99 | 0.975 |
| GCP-2^†^ | 177.8  (99.72-351.2) | 203.8  (144.0-335.6) | 173.7  (125.6-411.9) | 212.0  (135.5-296.1) | >0.99 | >0.99 | >0.99 | 0.7070 | 171.8  (117.1-221.2) | 271.0  (134.0-407.3) | 223.1  (116.5-314.4) | 0.2209 | 0.3526 | >0.99 | 0.990 |
| CXCL11^†^ | 58.34  (43.09-125.5) | 359.2  (186.4-519.7) | 143.0  (90.89-168.3) | 92.39  (83.09-119.8) | 0.0806 | **<0.0001** | **0.0047** | 0.5489 | 365.9  (96.34-676.7) | 118.7  (26.66-248.8) | 58.46  (17.84-237.7) | **0.0219** | **0.0024** | >0.99 | 0.990 |
| MIP-1β^†^ | 256.8  (191.9-305.6) | 357.9  (306.3-404.8) | 308.4  (279.6-403.3) | 276.6  (231.3-339.2) | 0.6177 | **0.0170** | 0.3992 | 0.5489 | 308.9  (228.9-375.8) | 284.8  (199.7-396.1) | 256.8  (194.1-279.1) | 0.7907 | 0.3526 | >0.99 | 0.990 |
| CCL1^†^ | 3.50  (2.30-4.288) | 16.01  (8.01-42.45) | 8.40  (3.99-18.46) | 4.49  (3.68-7.140 | **0.0342** | **<0.0001** | **0.0080** | 0.1577 | 13.90  (10.16-18.87) | 8.38  (3.84-29.22) | 3.92  (1.8-5.29) | >0.99 | **0.0010** | **0.0052** | 0.990 |
| IP-10^†^ | 27.53  (19.74-98.70) | 224.9  (98.06-308.1) | 92.13  (44.46-200.0) | 48.03  (28.65-97.27) | **0.008** | **<0.0001** | **0.0342** | 0.4473 | 130.5  (83.74-315.9) | 85.18  (38.12-143.9) | 43.13  (10.83-109.5) | 0.5391 | **0.0219** | 0.5391 | 0.990 |
| OSM^†^ | 460.8  (319.5-537.5) | 521.1  (437.5-656.0) | 521.1  (399.3-880.0) | 491.0  (344.6-686.6) | >0.99 | 0.7070 | 0.1733 | 0.6349 | 460.9  (391.5-720.9) | 497.6  (356.2-584.6) | 491.5  (273.0-568.5) | 0.7907 | 0.3526 | >0.99 | 0.990 |
| IL-33R^§^ | 6.48  (4.43-8.9) | 17.25  (11.78-28.89) | 12.84  (9.12-20.89) | 12.58  (6.32-14.25) | **0.0027** | **0.0027** | >0.99 | 0.0874 | 33.78  (22.52-70.52) | 12.03  (9.10-20.54) | 9.58  (6.47-12.16) | 0.2209 | **0.0110** | 0.7907 | 0.878 |
| OPN^§^ | 14.83  (12.46-23.52) | 25.90  (11.05-41.96) | 20.30  (9.895-39.58) | 23.00  (8.563-35.00) | 0.2460 | >0.99 | 0.3415 | 0.5489 | 47.40  (23.11-79.40) | 23.51  (10.88-40.52) | 22.68  (14.22-42.77) | 0.0760 | 0.4383 | >0.99 | 0.872 |
| PDGF-BB^§^ | 3.49  (1.93-4.97) | 5.23  (2.34-8.74) | 3.05  (1.83-7.38) | 2.29  (1.45-4.67) | 0.3415 | **0.0047** | 0.3415 | 0.6425 | 4.94  (2.64-8.13) | 3.25  (2.64-7.00) | 2.13  (0.97-3.78) | >0.99 | **0.0417** | 0.3526 | 0.975 |
| TM^§^ | 0.03  (0.02-0.10) | 0.05  (0.02-0.14) | 0.09  (0.02-0.15) | 0.10  (0.04-0.12) | >0.99 | >0.99 | >0.99 | 0.1293 | 0.03  (0.02-0.07) | 0.06  (0.03-0.13) | 0.105  (0.02-0.15) | 0.1009 | 0.0760 | >0.99 | 0.753 |
| CD163^§^ | 614.5  (323.5-827.2) | 829.4  (428.5-1582) | 792.8  (362.8-1225) | 823.2  (393.7-1077) | >0.99 | 0.2460 | 0.8051 | 0.4473 | 547.1  (366.3-1053) | 660.0  (343.2-1153) | 501.5  (267.0-1061) | >0.99 | 0.7907 | 0.3526 | 0.990 |
| IL-6Rα^§^ | 29.51  (22.89-34.30) | 33.86  (28.81-38.72) | 32.42  (27.77-38.98) | 32.36  (31.06-37.61) | >0.99 | >0.99 | 0.8051 | 0.0874 | 26.45  (22.66-33.61) | 30.34  (26.10-37.54) | 31.99  (26.35-35.77) | >0.99 | 0.5391 | >0.99 | 0.975 |
| CD30^†^ | 64.01  (35.13-92.94) | 145.3  (110.4-268.5) | 107.5  (73.42-256.2) | 104.7  (53.07-145.9) | >0.99 | **0.0133** | 0.1733 | 0.5489 | 94.36  (64.56-140.8) | 60.01  (42.69-156.3) | 51.40  (35.32-72.80) | >0.99 | 0.1325 | 0.5391 | 0.975 |
| IL-2Rα^§^ | 0.33  (0.24-0.66) | 1.82  (1.13-2.40) | 0.82  (0.54-1.47) | 0.68  (0.42-0.90) | **0.0342** | **<0.0001** | **0.0342** | 0.0874 | 1.43  (1.17-2.11) | 0.53  (0.365-1.12) | 0.49  (0.26-0.74) | 0.0760 | **0.0004** | 0.3526 | 0.975 |
| ApoA-I^‡^ | 2050  (364.2-7118) | 5264  (175-14620) | 5098  (335.2-7264) | 5519  (454.2-7081) | >0.99 | >0.99 | >0.99 | 0.6464 | 204.3  (101-29955) | 7496  (461-11029) | 6120  (249.1-8976) | >0.99 | >0.99 | >0.99 | 0.975 |
| Apo CIII^‡^ | 185.5  (64.70-284.3) | 177.9  (126.7-234.4) | 146.5  (86.05-198.0) | 125.2  (102.6-172.5) | >0.99 | 0.4642 | >0.99 | 0.5489 | 114.6  (78.89-376.5) | 188.8  (108.3-323.1) | 167.1  (96.66-250.8) | 0.7907 | >0.99 | >0.99 | 0.990 |

Week 24 was further compared and LTBI. Statistical comparisons were performed using a Friedman test adjusted for multiple comparisons (Dunn’s test) for BL v W6/8, BL v W24 and W6/8 v W24 for both PCTB and PTB groups and the Mann-Whitney test to compare LTBI with W24 in both PCTB and PTB with p-values adjusted using the Bonferroni method.

^‡^: Values are shown in mg/L, ^†^: Values are shown in pg/mL, ^§^: Values are shown in ng/mL.

Abbreviations. Apo: Apolipoprotein, C3: Complement component 3, C4: Complement component 4, CRP: C reactive protein, SAP: Serum amyloid protein, IL: Interleukin, TM: Thrombomodulin, PDGF: Platelet-derived growth factor, MCP: monocyte chemotactic protein, ICAM: Intercellular adhesion molecule, NCAM: Neural cell adhesion molecule, IP-10: Interferon γ-induced protein 10 kDa, OPN: Osteopontin, CD: Cluster of differentiation, MIG: Monokine induced by interferon gamma, G-CSF: Granulocyte colony-stimulating factor, IFN-γ: Interferon-gamma, OSM: Oncostatin M, VEGF: Vascular endothelial growth factor, MIP: Macrophage inflammatory protein, GCP-2: Granulocyte chemotactic protein 2, CXCL: C-X-C motif ligand, CCL: C-C motif ligand, M-CSF: Macrophage colony-stimulating factor, TNF-⍺: Tumour necrosis factor alpha, and TGF-β: Transforming growth factor beta.

**Supplementary table 4. Comparing the performance of HLA-DR expression and biosignatures from literature in discriminating LTBI from TB disease.**

**Performance of HLA-DR expression and biosignatures from literature in discriminating LTBI from PTB.**

| **Biosignature** | **AUC** | **Sensitivity (95% CI)** | **Specificity (95% CI)** | **Source** |
| --- | --- | --- | --- | --- |
| HLA-DR on Mtb-sp CD4 cells | 0.97 | 94.74% (75.36 - 99.73) | 100% (78.47 - 100) | Current study |
| CCL1 + CRP | 0.98 | 65% (40.78 – 84.64) | 100% (79.41 – 100) | Mutavhatsindi et al. [1] |
| IL-6Rα + IL-2Rα | 0.95 | 75% (50.90 – 91.34) | 93.75% (69.77 – 99.84) | Eribo et al. [2] |
| TNF-α + IL-12p40 | 0.92 | 85% (62.11 – 96.79) | 93.75% (69.77 – 99.84) | Sutherland et al. [3] |
| CCL1 + TNF-α | 0.92 | 65% (40.78 – 84.61) | 93.75% (69.77 – 99.84) | Chendi et al. [4] |
| IFN-𝛾 + IL-10 + IL-12p40 | 0.84 | 70% (45.72 - 88.11) | 75% (46.62 - 92.73) | Sutherland et al. [3] |
| IL-12p40 + IL-10 | 0.72 | 55% (31.53 - 76.94) | 87.50% (61.65 - 98.45) | Sutherland et al. [3] |

**Performance of HLA-DR expression and biosignatures from literature in discriminating LTBI from PCTB.**

| **Biosignature** | **AUC** | **Sensitivity (95% CI)** | **Specificity (95% CI)** | **Source** |
| --- | --- | --- | --- | --- |
| HLA-DR on Mtb-sp CD4 cells | 0.94 | 93.75% (71.6 - 99.68) | 100% (78.47 - 100) | Current study |
| CCL1 + CRP | 1.00 | 83.33% (58.58 – 96.42) | 100% (79.41 – 100) | Mutavhatsindi et al. [1] |
| IL-6Rα + IL-2Rα | 0.96 | 72.22% (46.52 – 90.31) | 87.50% (61.65 – 98.45) | Eribo et al. [2] |
| IFN-𝛾 + IL-10 + IL-12p40 | 0.89 | 61.11% (35.75-82.70) | 93.75% (69.77 – 99.84) | Sutherland et al. [3] |
| CCL1 + TNF-α | 0.85 | 61.11% (35.75-82.70) | 93.75% (69.77 – 99.84) | Chendi et al. [4] |
| TNF-α + IL-12p40 | 0.84 | 77.78% (52.36 – 93.59) | 93.75% (69.77 – 99.84) | Sutherland et al. [3] |
| IL-12p40 + IL-10 | 0.64 | 77.78% (52.36 – 93.59) | 62.50% (35.43 – 84.80) | Sutherland et al. [3] |

**Performance of HLA-DR and biosignatures from literature in discriminating LTBI from any aTB (PTB + PCTB).**

| **Biosignature** | **AUC** | **Sensitivity (95% CI)** | **Specificity (95% CI)** | **Source** |
| --- | --- | --- | --- | --- |
| HLA-DR on Mtb-sp CD4 cells | 0.96 | 94.29% (81.39 - 98.98) | 100% (78.47 - 100) | Current study |
| CCL1 + CRP | 0.98 | 71.05% (54.10 – 84.58) | 100% (79.41 – 100) | Mutavhatsindi et al. [1] |
| IL-6Rα + IL-2Rα | 0.95 | 76.32% (59.76 – 88.56) | 93.75% (69.77 – 99.84) | Eribo et al. [2] |
| CCL1 + TNF-α | 0.89 | 60.53% (43.39 – 75.96) | 93.75% (69.77 – 99.84) | Chendi et al. [4] |
| TNF-α + IL-12p40 | 0.88 | 76.32% (59.76 – 88.56) | 93.75% (69.77 – 99.84) | Sutherland et al. [3] |
| IFN-𝛾 + IL-10 + IL-12p40 | 0.85 | 68.42% (51.35 – 82.50) | 87.50% (61.65 – 98.45) | Sutherland et al. [3] |
| IL-12p40 + IL-10 | 0.69 | 55.26% (38.30 – 71.38) | 62.50% (35.43 – 84.80) | Sutherland et al. [3] |

Abbreviations. CRP: C reactive protein, IL: Interleukin, IFN-γ: Interferon-gamma, CCL: C-C motif ligand, and TNF-⍺: Tumour necrosis factor alpha.

**References**

[1]	H. Mutavhatsindi, G.D. Van Der Spuy, S. Malherbe, J.S. Sutherland, A. Geluk, H. Mayanja Kizza, A.C. Crampin, D. Kassa, R. Howe, A. Mihret, J.A. Sheehama, E. Nepolo, G. Günther, H.M. Dockrell, P.L. Lam Corstjens, K. Stanley, G. Walzl, N.N. Chegou, Validation and optimisation of host immunological bio-signatures for a point-of-care test for TB disease, Front. Immunol. 12 (2021). https://doi.org/10.3389/fimmu.2021.607827.

[2]	O.A. Eribo, M.S. Leqheka, S.T. Malherbe, S. McAnda, K. Stanley, G.D. van der Spuy, G. Walzl, N.N. Chegou, Host urine immunological biomarkers as potential candidates for the diagnosis of tuberculosis, Int. J. Infect. Dis. 99 (2020) 473–481. https://doi.org/10.1016/j.ijid.2020.08.019.

[3]	J.S. Sutherland, B.C. de Jong, D.J. Jeffries, I.M. Adetifa, M.O.C. Ota, Production of TNF-α, IL-12(p40) and IL-17 Can Discriminate between Active TB Disease and Latent Infection in a West African Cohort, PLOS ONE. 5 (2010) e12365. https://doi.org/10.1371/journal.pone.0012365.

[4]	B.H. Chendi, H. Tveiten, C.I. Snyders, K. Tonby, S. Jenum, S.D. Nielsen, M. Hove-Skovsgaard, G. Walzl, N.N. Chegou, A.M. Dyrhol-Riise, CCL1 and IL-2Ra differentiate Tuberculosis disease from latent infection Irrespective of HIV infection in low TB burden countries, J. Infect. 83 (2021) 433–443. https://doi.org/10.1016/j.jinf.2021.07.036.
